# A rise in double-strand breaks sensitizes tumours to oxidative metabolism inhibitors

**DOI:** 10.1101/2023.12.19.572355

**Authors:** Ferran Medina-Jover, Agnès Figueras, Álvaro Lahiguera, Pau Guillén, Roderic Espín, Miguel Ángel Pardo, Miquel Angel Pujana, Edurne Berra, Alberto Villanueva, Adrià Bernat-Peguera, Margarita Romeo, José Carlos Perales, Francesc Viñals

**Affiliations:** Departament de Ciències Fisiològiques, Universitat de Barcelona, 08908 L’Hospitalet de Llobregat, Barcelona, Spain; Program Against Cancer Therapeutic Resistance (ProCURE), Institut Català d’Oncologia (ICO), Hospital Duran i Reynals, 08908 L’Hospitalet de Llobregat, Barcelona, Spain; Oncobell Program, Institut d’Investigació Biomèdica de Bellvitge (IDIBELL), 08908 L’Hospitalet de Llobregat, Barcelona, Spain; Servei d’Oncologia Mèdica, Institut Català d’Oncologia (ICO) Hospital Duran i Reynals, 08908 L’Hospitalet de Llobregat, Barcelona, Spain; Center for Cooperative Research in Biosciences (CIC bioGUNE), Basque Research and Technology Alliance (BRTA), Bizkaia Technology Park, Building 801A, 48160, Derio, Spain; CIBERONC, Madrid, Spain; Badalona Applied Research Group in Oncology (B-ARGO), Institut d’Investigació en Ciències de la Salut Germans Trias i Pujol (IGTP), 08916 Badalona, Spain; Medical Oncology Department, Institut Català d’Oncologia Badalona, Badalona Applied Research Group in Oncology (B-ARGO), Institut d’Investigació Germans Trias i Pujol (IGTP), Carretera del Canyet s/n, 08916 Badalona, Spain

## Abstract

Double strand brakes (DSB) accumulate in cellular DNA as a result of deficiencies in homologous recombination repair systems, such as mutations in *BRCA* genes, or upon antitumoral treatments. In the present study we show that the accumulation of DSB, regardless of its origins, leads to a shift towards oxidative metabolism. We have identified that DSB-induced reactive oxygen species (ROS) promote the activation of NRF2 which downregulates the glycolytic transcription factor HIF-1. HIF-1 inhibition is a key step in this metabolic shift, because leads to the reduction of PDHK1 levels and the consequential activation of pyruvate dehydrogenase, a mitochondrial gatekeeper of cellular metabolism, promoting this metabolic shift. Remarkably, after the induction of DSBs, the tumour is more sensitive to the inhibition of oxidative metabolism since both treatments synergize in vivo, resulting in reduced tumour growth. Therefore, we demonstrate a significant feedback between DSBs induction and cancer cell metabolism that ultimately limits the cell’s potential for metabolic plasticity, hence sensitizing it to the action of counteracting drugs.

## INTRODUCTION

Alterations in DNA repair are a hallmark in the tumorigenic process. Repairing double strand brakes (DSBs) is an utmost critical task to cope with potential for compromising chromosome integrity^1,2^.DSB repair involves three different mechanisms: homologous recombination pathway (HR), non-homologous end joining (NHEJ), and alternative end joining (A-EJ). In the tumoral context, the HR pathway is crucial. Mutations in genes of these pathways increase genomic instability of the tumour^3,4^. Specially, mutations in genes related to HR, such in *BRCA1/2* or *RAD51* but also epigenomic silencing, have been broadly described in diverse tumour types, being of great relevance in ovarian, breast and prostate cancer^4–6^. This type of tumours (HR-deficient, HRD) become increasingly reliant on A-EJ, particularly on the activity of PARP1^7–9^, which detects DNA damage and undergoes poly-ADPribosylation (PAR) along with other proteins involved in the DNA damage response^10^.

Inducing DNA damage is a common mechanism of action for several chemotherapeutic agents used in the clinics. For instance, DSB promoting chemotherapeutics inhibit topoisomerases or the synthesis of nucleotides. In the case of ovarian cancer, these chemotherapies are used when patients develop resistance to platinum-based therapies. However, in other tumour types (i.e. breast cancer) they are the first line of treatment. Regarding tumours with HRD, they respond better to classical chemotherapies due to its reduced capacity to repair DNA^11^. Further, in the last decade, the clinical effectiveness of PARP inhibitors (PARPi) has also been well-established for HRD tumours, not only in ovarian cancer but also in breast and prostate cancer^12–20^. The use of PARPi has been proved highly beneficial in managing these types of cancers in the medical setting.

NAD^+^ is a critical factor in PARP activity, as it acts as the substrate for poly- ADPribosilation. In addition, NAD^+^ serves as a vital coenzyme in numerous reactions within both glycolysis and the Krebs cycle. During these processes, NAD^+^ is reduced to NADH, which subsequently donates its electrons to another acceptor to regenerate NAD^+^. In normal, non-transformed cells, NADH predominantly transfers electrons to the respiratory chain, generating substantial amounts of ATP. However, in tumoral cells exhibiting the Warburg phenotype, NADH donates its electrons to pyruvate, resulting in the production of lactate, which is then secreted. We have previously described that HRD tumours display a more oxidative metabolism compared to tumours with HR proficiency^21^.HRD tumours rely on PARP1 for DNA repair, a process that demands significant amounts of NAD^+^ to be active. Oxidative metabolism in these tumours enables maintained PARP1 activity, as the respiratory chain facilitates a faster regeneration of NAD^+^. Mechanistically, HRD tumours enhance OXPHOS (oxidative phosphorylation) metabolism by increasing the carbon flux, derived from glucose, into the Krebs cycle. This flux regulation is largely governed by the pyruvate dehydrogenase (PDH) enzyme, responsible for converting pyruvate into the first substrate of the Krebs cycle, acetyl-CoA. PDH regulation is complex, involving different allosteric regulations and three phosphorylation sites that can inhibit its activity^22^. Notably, the phosphorylation of Serine-293 shows the strongest correlation with PDH activity^23^. Pyruvate dehydrogenase kinases (PDHK) are responsible for phosphorylating these sites, with four isoforms described, which have different affinities for each residue^24^.

In this study, we unveil an intriguing connection between DSB and the shift towards an oxidative metabolism. Specifically, DNA damage triggers the downregulation of glycolytic transcription factors that govern PDH activity. Furthermore, by combining DSB inducers, like doxorubicin, with respiratory chain inhibitors, we observe a remarkable increase in tumour cell death across various cancer types. This therapeutic approach has the potential to significantly enhance the effectiveness of current treatments, making it particularly valuable in managing relapse, a common clinical scenario in HGSOC patients.

## METHODS

### Bioinformatics

The Cancer Genome Atlas (TCGA) data were downloaded from GDC data portal (https://portal.gdc.cancer.gov/). RNA-seq data was used to computed scores using computational single sample gene set enrichment analysis (ssGSEA) method^25^ for the gene signatures Oxidative phosphorylation (KEGG pathway hsa00190), MOOTHA_VOXPHOS^26^, respiratory chain complex I (GO:0045279), HIF1-specific targets (KEGG pathway hsa04066) and a custom gene signature derived from DDR-HDR pathway definition^27^. Pearson correlation coefficients were estimated between gene signatures for each tumour type using cor function, and p-value and confidence interval were calculated using cor.test, both functions from R package stats version 4.3.1.

Cell lines sensitivity data to dihydrorotenone were downloaded from Sanger Institute (https://www.cancerrxgene.org/). RMA normalised basal expression profiles for these cell lines were also downloaded from this database. Scores calculated using ssGSEA method^25^ were correlated with IC50 sensitivity to dihydrorotenone values using Spearman’s rank correlation coefficient. ssGSEA were selected from the canonical pathways of the curated gene sets of the GSEA and were related to DNA damage, glycolytic metabolism or oxidative metabolism. Genesets are shown if Spearman estimate is >|0.1| and FDR<0.05. From R package stats version 4.3.1, cor function was used for coefficient calculation, and cor.test for p-value calculation.

For the ChIP analysis, data was obtained from ChIP-Atlas^28^ using the Peak Browser tool and later visualized with the Integrative Genome Viewer (IGV)^29^.

### Cell Culture

The ID8 cell lines were kindly donated by Dr Iain McNeish (Imperial College, London, UK) and have been described previously^30^. Cells were cultured in Dulbecco’s modified Eagle’s medium (DMEM, 25 mM glucose, 1 mM pyruvate, Biowest) supplemented with 4% foetal bovine serum (FBS), 5mM L-glutamine, ITS (5 μg/ml insulin, 5 μg/ml transferrin, and 5 ng/ml sodium selenite, Gibco), and antibiotics (100 U/ml penicillin and 0.1 mg/ml streptomycin). SKOV3N cells were purchased from Sigma-Aldrich. Cells were cultured in DMEM (25 mM glucose, 1 mM pyruvate, Sigma-Aldrich) supplemented with 10% FBS, 5mM L-glutamine, and antibiotics (100 U/ml penicillin and 0.1 mg/ml streptomycin). OVCAR4 cells were kindly donated by Dr Hamilton (FoxChase Cancer Center, Philadelphia, U.S.) and were grown in RPMI 1640 Medium (Gibco, Cat. Num. #21875034) supplemented with 10% FBS, 2mM L-glutamine, 1mM pyruvate, antibiotics and 0.25 units/mL Insulin. A2780 cells were kindly donated by Dr Josefa Giménez-Bonafé (Universitat de Barcelona, Barcelona, Spain). Cells were grown in RPMI 1640 Medium supplemented with 10% FBS, 10mM L-glutamine, and antibiotics (100 U/ml penicillin and 0.1 mg/ml streptomycin). NP P53, 22Rv1 and DU145 cells were kindly donated by Dr Alvaro Aytés (ProCure Program, Institut Català d’Oncologia, Hospitalet de Llobregat, Spain) and were grown in RPMI Medium (Gibco) supplemented with 10% FBS, 5mM L-glutamine, and antibiotics (100 U/ml penicillin and 0.1 mg/ml streptomycin). T-47D cells were kindly donated by Dr Miquel Angel Pujana (ProCure Program, Institut Català d’Oncologia, Hospitalet de Llobregat, Spain) and were grown with RPMI Medium (Gibco) supplemented with 10% FBS, 10mM L-glutamine, and antibiotics (100 U/ml penicillin and 0.1 mg/ml streptomycin).

### Cell death assay

Cells were seeded in 12-well plates. The day after, cells were treated with the respective compounds and solvents for 24 hours. Then cells were detached from the plate using TrypLE dissociation reagent (Gibco) and inactivated with 2mL of complete media. Cells were centrifuged at 1000g for 5 minutes and washed with PBS 3 times. Next, cells were resuspended with 200 µL of PBS with 1 µg/mL of propidium iodide (Sigma-Aldrich). Flow cytometry was performed in a Gallios flow cytometer and data analysed with Kaluza software (Beckman Coulter, Brea, CA, USA).

### Cell viability assay

Cells well were seeded in 96-well plates, doing 8 replicates. After attachment, cells were treated with the respective compounds and solvents for 72 hours. Then cells were fixed with methanol for 10 minutes, and then stained for 20 minutes with Crystal Violet (0.1% in ddH2O). Cells were washed 3 times with water and then crystal violet was resolubilized (5% acetic acid, 10% methanol in ddH2O) and optic density was measured with a multiwell plate reader at 595 nm. For synergy assays the same protocol was used, but the analysis was performed using the Combenefit software^31^.

### Immunohistochemistry in paraffin embedded samples and scoring

Paraffin-embedded sections were deparaffinized in xylene and rehydrated in downgraded alcohols and distilled water. Antigen retrieval was performed in a decloaking chamber (Decloaking Chamber™ NxGen, Bio Care), using Dako retrieval buffers, depending on the antibody used (HIF-1α, Dako Target Retrieval Solution pH 6; for BACH-1, Dako Target Retrieval Solution pH 9). Next, endogen peroxidases were deactivated (3% H_2_O_2_, 30% Methanol in water) for 15 minutes and then permeabilized with PBS-T (PBS, 0.1% Triton 100X). Samples were blocked with goat serum diluted 1:200 in PBS for one hour followed by an overnight incubation at 4°C with the primary antibody (HIF-1α, 1:200, described in^32^; BACH-1, 1:200, Sigma Aldrich HPA034949) Next, samples were washed 3 times with PBS-T, and incubated at room temperature for an hour with Envision Dual Link (Dako) followed by three washes with PBS-T. After the DAB+ developing system (Dako) was used to stain the samples followed by a counterstain with haematoxylin and visualization with a microscope. For scoring, several representative pictures were taken from each preparation and then analysed with H-Score tool from QuPath^33^. All patient samples were processed following standard operating procedures, with the appropriate approval of the Clinical Investigation Ethics Committee of HUB (meeting minutes 18/11, ref. PR236/11) and conformed to the principles set out in the WMA Declaration of Helsinki. A signed informed consent was obtained from each patient.

### In-vivo experiments

2 million ID8-P53 cells were injected orthotopically in the right ovary of 8 to 10 weeks old C57BL6J female mice. After one month, mice were randomized in 4 different groups. Liposomal doxorubicin (Caelyx) was bought from Accord Healthcare S.L.U and prepared by the Pharmacy Department of the Institut Català d’Oncologia at 2mg/mL. Just before administration, Caelyx was diluted in saline 5% glucose to a final concentration of 0.5 mg/mL. IACS-10759 (MedChemExpress, HY-112037) was dissolved in 0.5% carboxymethyl cellulose at 37.5 µg/mL. Mice were treated for 4 weeks with 2mg/kg of Caelyx which was given once a week with an intraperitoneal injection, and 0.3 mg/kg of IACS was given 5 times a week by oral gavage. Twice a week each animal received a subcutaneous injection of 300µL of saline 5% glucose.

Weight was measured once a week. For glucose and lactate levels, blood was obtained from the saphenous vein. Glucose was measured with reactive strips using Glucomen® machine (Menarini Diagnostics), and lactate was measured with reactive strips using Laktate machine (Eaglenos, Busimedic, S.L.). Glucose and lactate levels were measured every 2 weeks. Health and wellbeing of the animals was strictly monitored, and they were euthanized when endpoint criteria was met. None of administered treatments nor mice manipulation had a significative effect on mice body weight and behaviour. All studies were approved by the local committee for animal care (IDIBELL, DAAM 5766)

### Lentiviral production and infection

3 million of HEK-293FT cells were seeded in a 100 mm cell culture dish with DMEM (25 mM glucose, 1 mM pyruvate, Sigma-Aldrich) supplemented with 10% FBS and 5mM L-glutamine, and antibiotics (100 U/ml penicillin and 0.1 mg/ml streptomycin). The following day, cells were transfected with 7.5 μg of psPAX (Addgene, plasmid # 12260), 5 μg of pVSV-G (Addgene, plasmid # 8454), and 10 μg of the desired plasmid, diluted in a final volume of 500 μL of Optimem (Gibco). Next, this volume was mixed 1:1 with a solution of polyetylenimide (PEI, Sigma) in Optimem, following manufacturer’s instructions, and then incubated at room temperature for 30 minutes. Then, 1 mL of the mix was added in each dish mixed with 9 mL of basal DMEM for 7 hours. Later, cells were washed one time with PBS and then DMEM 1% FBS was added.

After 2 days, supernatants were filtered with 0.45 µM filters. Then 1mL of the supernatants was mixed with a 1mL of the desired cell line at a concentration of 200,000 cells/mL in a well of a 6-well plate. Then 8 µg/mL of polybrene was added, and immediately centrifuged at 32°C for 90 minutes at 1000g. Selection was started the day after.

ID8-P53 cells were infected, using CRISPR/CAS9 technology, with an empty vector (PC) or a vector against *Bach1* with the following gRNA 5’ GCGGTTCCGAGCCCACCGCT 3’ (GenScript Biotech Corporation, USA) (BACH1 KO) and selected using Bleomycin (10µg/mL, Merck). SKOV3N cells with *BRCA2* KO were generated and described previously^21^.

ID8-P53 cells were infected, using shRNA technology with and empty vector containing GFP gene or a vector with and shRNA against *Hif1a* gene (Horizon Discovery, UK) (5’ GCATTAAAGCAGCGTATC 3’) also containing GFP gene. Selection was performed using FACS, selecting positive cells for GFP.

ID8-P53 and ID8-BRCA cells were infected, using shRNA technology with and empty vector containing Tdtomato gene or a vector with and shRNA against *Nfe2l2* gene (Horizon Discovery, UK) (5’ CCCGAATTACAGTGTCTTAAT 3’) also containing Tdtomato gene. Selection was performed using FACS, selecting positive cells for Tdtomato.

### Quantitative real time PCR

RNA extraction was performed using TRIzol reagent (ThermoFischer Scientific), according to manufacturer’s instructions. cDNA was obtained from RNA using a reverse transcriptase (High-Capacity cDNA Reverse Transcription Kit; Applied Biosystems, Life Technologies, CA). qPCR was performed using cDNA in a Lightcycler instrument (Roche Molecular Biochemicals) using SybrGreen (Roche Molecular Biochemicals) according to manufacturer’s instructions. Primers sequences are described in the following table:

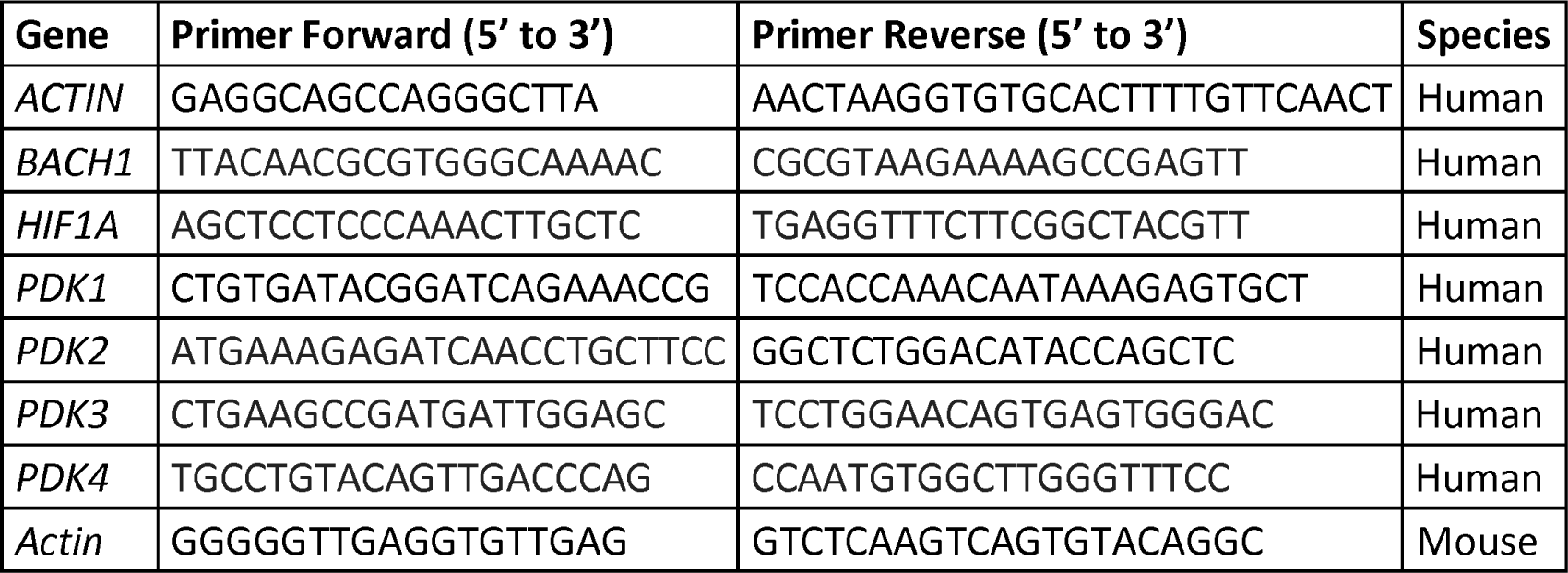
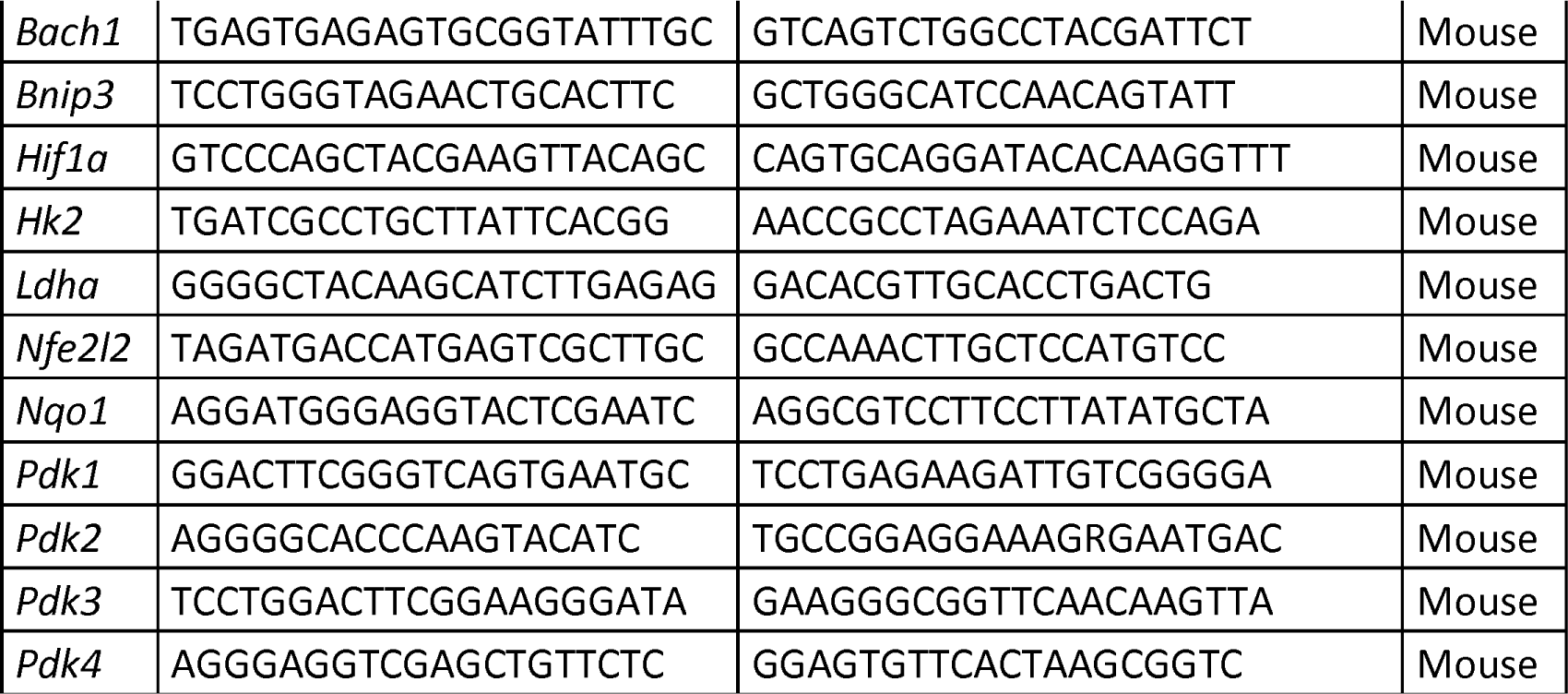

### Reagents

Other reagents that are not specified in the following table where purchased to Sigma- Aldrich:

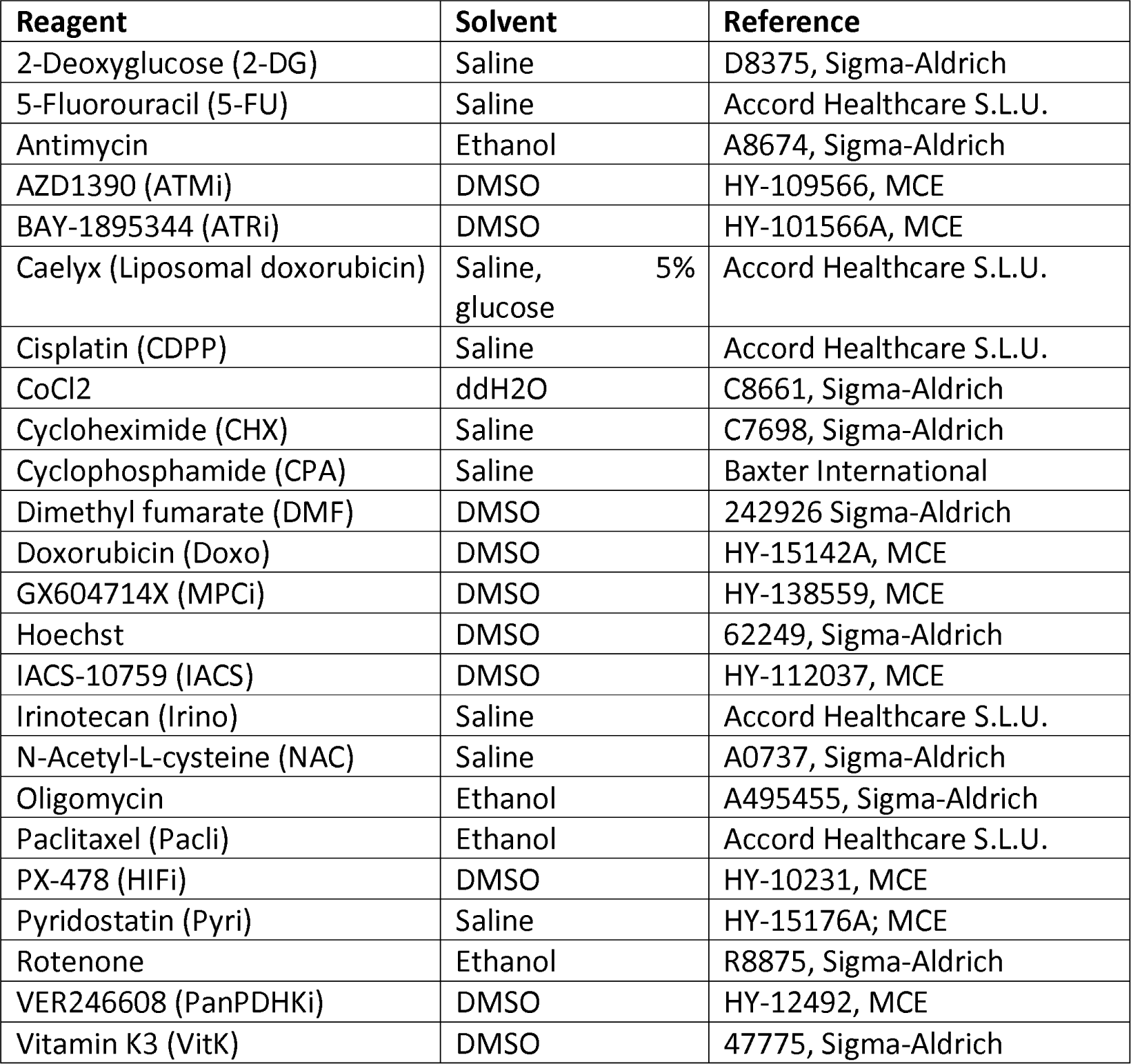

Reagents Other reagents that are not specified in the following table were purchased from Sigma-Aldrich:
| Reagent | Solvent | Reference |
| --- | --- | --- |
| 2-Deoxyglucose (2-DG) | Saline | D8375, Sigma-Aldrich |
| 5-Fluorouracil (5-FU) | Saline | Accord Healthcare S.L.U. |
| Antimycin | Ethanol | A8674, Sigma-Aldrich |
| AZD1390 (ATMi) | DMSO | HY-109566, MCE |
| BAY-1895344 (ATRi) | DMSO | HY-101566A, MCE |
| Caelyx (Liposomal doxorubicin) | Saline, 5% glucose | Accord Healthcare S.L.U. |
| Cisplatin (CDPP) | Saline | Accord Healthcare S.L.U. |
| CoCl<sub>2</sub> | ddH<sub>2</sub>O | C8661, Sigma-Aldrich |
| Cycloheximide (CHX) | Saline | C7698, Sigma-Aldrich |
| Cyclophosphamide (CPA) | Saline | Baxter International |
| Dimethyl fumarate (DMF) | DMSO | 242926 Sigma-Aldrich |
| Doxorubicin (Doxo) | DMSO | HY-15142A, MCE |
| GX604714X (MPCi) | DMSO | HY-138559, MCE |
| Hoechst | DMSO | 62249, Sigma-Aldrich |
| IACS-10759 (IACS) | DMSO | HY-112037, MCE |
| Irinotecan (Irino) | Saline | Accord Healthcare S.L.U. |
| N-Acetyl-L-cysteine (NAC) | Saline | A0737, Sigma-Aldrich |
| Oligomycin | Ethanol | A495455, Sigma-Aldrich |
| Paclitaxel (Pacli) | Ethanol | Accord Healthcare S.L.U. |
| PX-478 (HIFi) | DMSO | HY-10231, MCE |
| Pyridostatin (Pyri) | Saline | HY-15176A; MCE |
| Rotenone | Ethanol | R8875, Sigma-Aldrich |
| VER246608 (PanPDHKi) | DMSO | HY-12492, MCE |
| Vitamin K3 (VitK) | DMSO | 47775, Sigma-Aldrich |

### ROS measurement

Cells were seeded in 12-well plates. The day after, cells were treated with the respective compounds and solvents for 24 hours. Then cells were detached from the plate using TrypLE dissociation reagent (Gibco) and inactivated with 2mL of complete media. Cells were centrifuged at 1000g for 5 minutes and washed with PBS 3 times. Next, cells were resuspended with 100µL of CM-H2DCFA (Invitrogene, Ref #C6827) 2.5 µM in PBS and incubated at 37°C in the dark for 45 minutes. Then 2mL of PBS was added and cells were centrifuged at 1000g for 5 minutes followed by 2 washes with PBS. Finally, cells were resuspended with 200µL PBS with 1 µg/mL of propidium iodide (Sigma-Aldrich). Flow cytometry was performed in a Gallios flow cytometer and data analysed with Kaluza software (Beckman Coulter, Brea, CA, USA).

### Seahorse Assay for Glycolitic flux and OCR assessment

Firstly, cells were seeded in a 96 well Seahorse (Agilent, Technologies Inc., Santa Clara, USA) plate in 80µL. Next day, 80µL of medium was added to each well containing the correspondent solvent or drug. The following day the Glycolytic flux assay was performed following the manufacturer’s instructions.

Normalization was performed using nuclei quantification. 2µM of Hoechst was added to each well, when the assay was finished, and the plate was incubated for 15 minutes at 37°C. Then, using the Zeiss Axio Observer Z1+ Apotome inverted fluorescence microscope, 9 images of each well were taken, accounting for the 40% of the well area. The nuclei were counted using Fiji software.

### Western Blot

For obtaining proteins from cells, cells were collected and washed twice in cold PBS and lysed for 20 min at 4 °C in RIPA lysis buffer (PBS pH 7.4, SDS 0.1%, NP-40 1%, sodium deoxycholate 0.5%) with protease and phosphatase inhibitors (100 μM PMSF, 40 mM β-glycerol phosphate, 1 μg/mL leupeptin, 4 μg/mL aprotinin, 1 μM pepstatin A, 0.1 μg/mL benzamidine, 200 μM sodium orthovanadate, 10 mM NaF, and 5 mM EDTA). For tissues, the sample was manually disaggregated with the use of liquid nitrogen. The material was collected and lysed with RIPA lysis buffer with protease and phosphatase inhibitors. Next, insoluble material was removed by centrifugation at 8000 g for 15 min at 4 °C. SDS/PAGE was used to separate proteins, which were then electrophoretically transferred to Immobilon-P membranes (Millipore, Burlington, MA, USA; Cat. Num. #IPVH00010) in 25 mm Tris/HCl, 190 mm glycine and 10% methanol at 4 °C. Membranes were blocked in TBS (10 mm Tris/HCl, 150 mm NaCl, pH 7.4) with 5% of non-fat dry milk at room temperature for 1 h. Blots were incubated with the corresponding antibody (see table below) in TBS 1% non-fat dry milk overnight. Next, blots were washed in TBS 0.1% Triton X-100 followed by an incubation with anti-rabbit Ig or anti-mouse (Amersham Pharmacia Biotech, Little Chalfont, UK; Cat. Num. #NA934 and #NXA931, respectively) horseradish peroxidase-linked antibodies, in TBS 1% non-fat milk. Blots were washed in TBS 0.1% Triton X-100. Then, ECL Prime (Amersham Pharmacia Biotech; Cat. Num. #RPN2236) was added and measured with the ChemiDoc™ MP imaging system. Volumetric analysis was performed using image lab from Bio-Rad (Hercules, CA, USA).

| **Protein** | **Dilution** | **Species** | **Reference** |
| --- | --- | --- | --- |
| Actin | 1:2500 | Mouse | Sigma Aldrich AC-15 A5441 |
| BACH-1 (for WB) | 1:1000 | Mouse | Santa Cruz Biotech sc-271211 |
| HIF-1α | 1:500 | Rabbit | Described in <sup>34</sup> |
| NRF2 | 1:500 | Rabbit | Cell Signaling #12721 |
| PDHK1 | 1:1000 | Rabbit | Cell Signaling #3820 |
| PDHK4 | 1:1000 | Rabbit | Novus Biologicals #NBP1-07049 |
| PDHT | 1:1000 | Rabbit | Cell Signaling #3205 |
| p-PDH Ser232 | 1:1000 | Rabbit | Sigma-Aldrich (AP1060) |
| p-PDH Ser293 | 1:1000 | Rabbit | Abcam (ab92696) |
| Tubulin | 1:2000 | Mouse | Sigma Aldrich T6079 |
| Vinculin | 1:2000 | Mouse | Sigma Aldrich V9131 |
| γ-H2AX | 1:500 | Rabbit | Cell Signaling #9718 |

### Statistical Analysis

Statistical analyses were performed with GraphPad Prism (version 8.3). Unless otherwise specified, data are presented as Mean ± SD, and each graph dot represents an independent biological replicate. Significant differences were assessed using Mann-Whitney U-test when comparing between two groups, and significant differences were considered when p-value<0.05.

## RESULTS

### DNA damage correlates with expression of OXPHOS related pathways

Our group previously described^21^, that chronic DNA damage resulting from a deficient HR system led to an increase in oxidative metabolism in breast and ovarian cancers which represented a therapeutic opportunity for those cancers. In order to demonstrate that is a generalised effect in cancer, we calculated the correlation between the HR deficiency index using the Knijnenburg HRD score^27^ and the expression of different OXPHOS signatures, using available genomic and mRNA data from the TCGA (Fig. 1A). We observed that in general, high HR deficiency correlated with higher expression of OXPHOS pathways across different tumour types such prostate, lung or pancreatic cancer. Next, using the Database of genomics of drug sensitivity in cancer of the Sanger Institute, we correlated the sensitivity of each cell line to Dihydrorotenone, an inhibitor of the complex I of the respiratory chain, and the value of the single-sample gene set expression analysis (ssGSEA) that were on the canonical pathways of the curated gene sets of the GSEA and related to DNA damage, glycolytic metabolism or oxidative metabolism (Fig. 1B, Sup. Table 1). As expected, cells with high expression of OXPHOS metabolism-related genes present a greater sensitivity to Dihydroreotenone and, on the contrary, those with higher expression of hypoxia or glycolysis were more resistant. Interestingly, gene expression related to DNA repair negatively correlated with resistance to Dihydrorotenone, suggesting that functional DNA damage repair pathways are dependent on oxidative metabolism.

**Figure 1.**
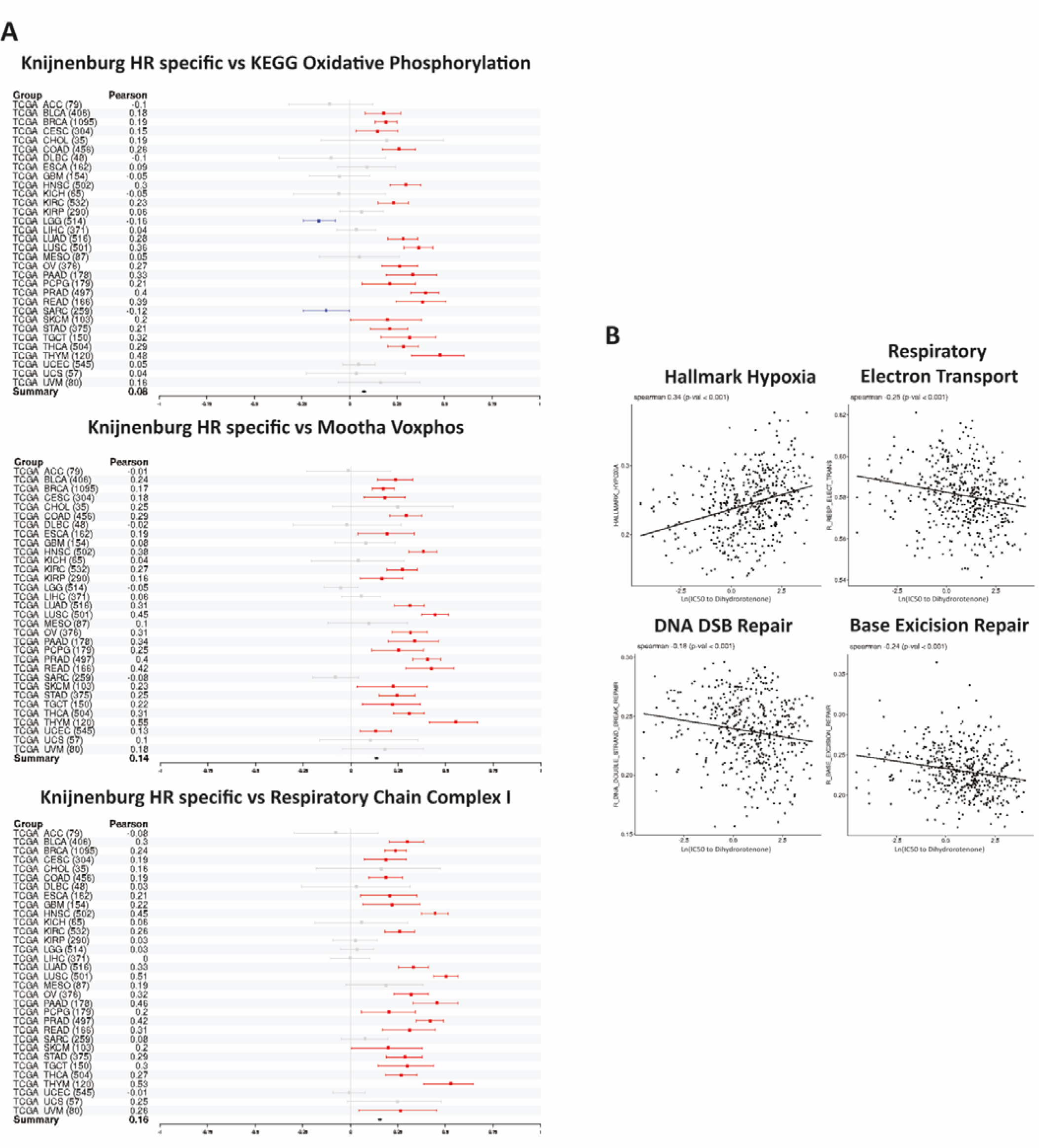
DNA damage correlates with an increased oxidative metabolism. A) Forest plots showing the correlation between Knijnenburg index and KEGG oxphos, Mootha voxphos and respiratory chain complex I signatures respectively in different cancer types (TCGA Study Abbreviations, Sup. Table 3). Red lines indicate significative positive correlation meanwhile blue indicates significative negative correlation. Grey indicates no significant correlation. **B)** Correlation between the sensitivity to dihydrorotenone, expressed as Ln (IC50), and Hallmark hypoxia, Respiratory electron transport, DNA DSB repair and Base excision repair, respectively, in different cell lines annotated as carcinoma, sarcoma or melanoma.

### DSB induction promotes oxidative metabolism

To validate our *in-silico* findings we examined if inducing acute DNA damage would also trigger a rise in oxidative metabolism. We measured the oxygen consumption rate (OCR) and the glycolysis rate in a HR proficient model: murine ovarian cancer ID8-KO *Trp53* cells (referred hereafter as ID8-P cells). These cells were treated with various chemotherapy agents used in clinical settings (Fig. 2A-B, panel left). Treatments included doxorubicin and irinotecan (topoisomerase inhibitors), 5-fluorouracil (thymidylate synthase inhibitor), paclitaxel (microtubule stabilizing agent) and cyclophosphamide and cisplatin (alkylating agents). Notably, only when promoting DSBs, measured as the presence of y-H2AX, our ID8-P model did exhibit a shift in the carbon flux towards an increased oxidative metabolism, measured as an increase in the OCR/glycolysis ratio. This was similar to what we observed in ID8 HR-deficient cells (KO for *Trp53* and *Brca2* genes, referred hereafter as ID8-B cells) which sustain chronic DNA damage due to the accumulation of DSB (Fig. 2A and B, panel right). In contrast, treatment with chemotherapy that did not induce DSBs (e.g Paclitaxel, CPA 0.1mM, CDPP 5 μM) did not cause an oxidative shift. However, these treatments did increase both the glycolytic flux and the OCR (Sup Fig. 1A and B). It is worth noting that very high doses of alkylating agents, which ultimately induce DSB (e.g. CPA 2mM, CDDP 40 μM), did eventually trigger the shift towards an increased oxidative metabolism (Fig. 2A). Remarkably, we did not observe an increased cell death in any treatment (Sup Fig 1C). Further, we observed an increased OCR/glycolysis ratio in different HR-proficient cell line models of ovarian, prostate and breast cancer (Fig 2C) when DSB where induced (Fig 2D). Overall, these data confirmed that the shift towards and OXPHOS metabolism when DSB are produced is a response mechanism shared across different cancer types.

**Figure 2.**
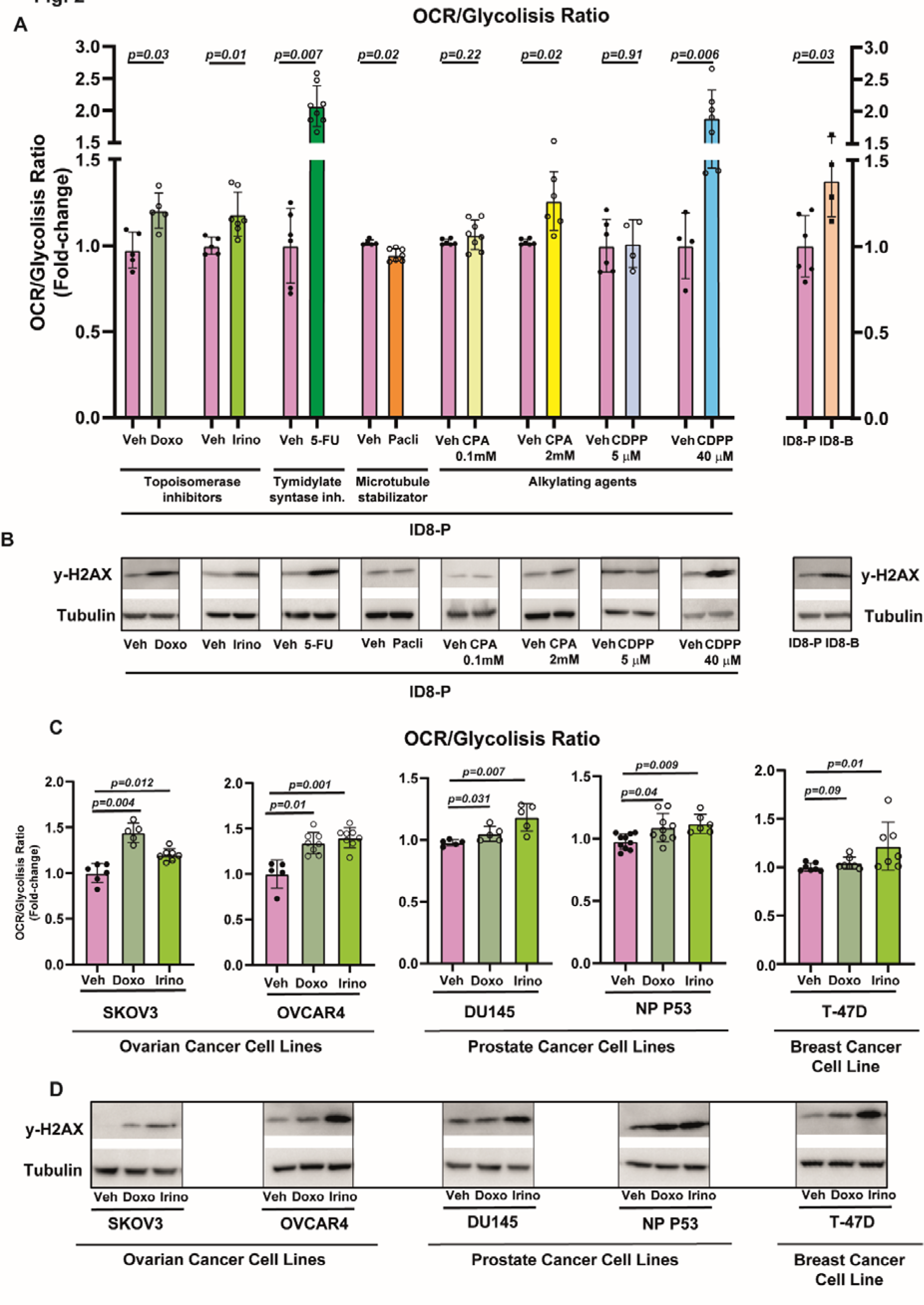
Acute DNA damage produces an increase in oxidative metabolism. A) OCR/Glycolysis ratio is shown in ID8-*Trp53* KO cells (ID8-P) pretreated for 24h with vehicle (Veh, DMSO), doxorubicin 2µM (Doxo), irinotecan 50µM (Irino), 5-fluorouracil 5µM (5-FU), paclitaxel 20nM (Pacli), ciclophosphamide 0.5mM or 2mM (CPA) or cisplatin 5µM or 40µM (CDPP) (left panel), and in ID8-P and ID8-*Trp53 Brca2* KO cells (ID8-B) (right panel). **B)** Western blot assessing the presence of y-H2AX in ID8-*Trp53* KO cells (ID8-P) pretreated for 24h with vehicle (Veh, DMSO), doxorubicin 2µM (Doxo), irinotecan 50µM (Irino), 5-fluorouracil 5µM (5-FU), paclitaxel 20nM (Pacli), ciclophosphamide 0.5mM or 2mM (CPA) or cisplatin 5µM or 40µM (CDPP) (left panel), and in ID8-P and ID8-*Trp53 Brca2* KO cells (ID8-B) (right panel). **C)** OCR/Glycolysis ratio is shown in SKOV3, OVCAR4, DU145, NP P53 and T-47D cells. Cells were pretreated for 24 hours with vehicle (Veh, DMSO), 2µM doxorubicin (doxo) or with irinotecan (irino) 50µM for OVCAR4, T-47D and DU145 and 20µM for SKOV3 and NP P53 cells. **D)** Western blot assessing the presence of y-H2AX and tubulin (as a loading control) in SKOV3, OVCAR4, DU145, NP P53 and T-47D cells. Cells were pretreated for 24 hours with vehicle (Veh, DMSO), 2µM doxorubicin (doxo) or with irinotecan (irino) 50µM for OVCAR4, T-47D and DU145 and 20µM for SKOV3 and NP P53 cells. Data are presented Mean ± SD unless otherwise specified, and each graph dot represents an independent biological replicate. Significant differences were assessed using Mann-Whitney U-test when comparing between two groups and considered when P<0.05. The p value is indicated in each figure.

### The PDHK1/PDH axis is responsible for up-regulating OXPHOS after DSB induction

In a previous study^21^ we described that the regulation of the PDH activity had a major role in regulating the energetic metabolism in HRD tumours. Therefore, we examined if acute DSB-DNA damage produced by chemotherapy could also impact the phosphorylation of pyruvate dehydrogenase (PDH). We observed a reduction of the phosphorylation of PDH in ovarian cancer cell lines, when DSB where induced independently of the drug used (Fig 3A-C, Sup. Fig. 2A-C), or the cell line (Sup. Fig. 2D- F). Further, we observed the same reduction in phosphorylation of PDH in cell lines of prostate (Fig 3D-F), and breast cancer (Fig 3G-I) treated with DSB inductors. Finally, to verify that this effect was indeed caused by the generation of DSBs, we incubated cells with pyridostatin, a compound that stabilizes G-quadruplex in cells, which specifically generates DSBs^35^ (Fig. 3J). Remarkably, this compound also decreased the phosphorylation of PDH (Fig. 3K-L). Likewise, we also observed the reduction in the phosphorylation of PDH after treating cells with (6 Gy) radiotherapy (Sup. Fig. 2G-H). All these results indicate that DSB generated either by chronic DNA damage due to HRD phenotype or acute DNA damage, which enhances OXPHOS metabolism, correlated with increased PDH activity.

**Figure 3.**
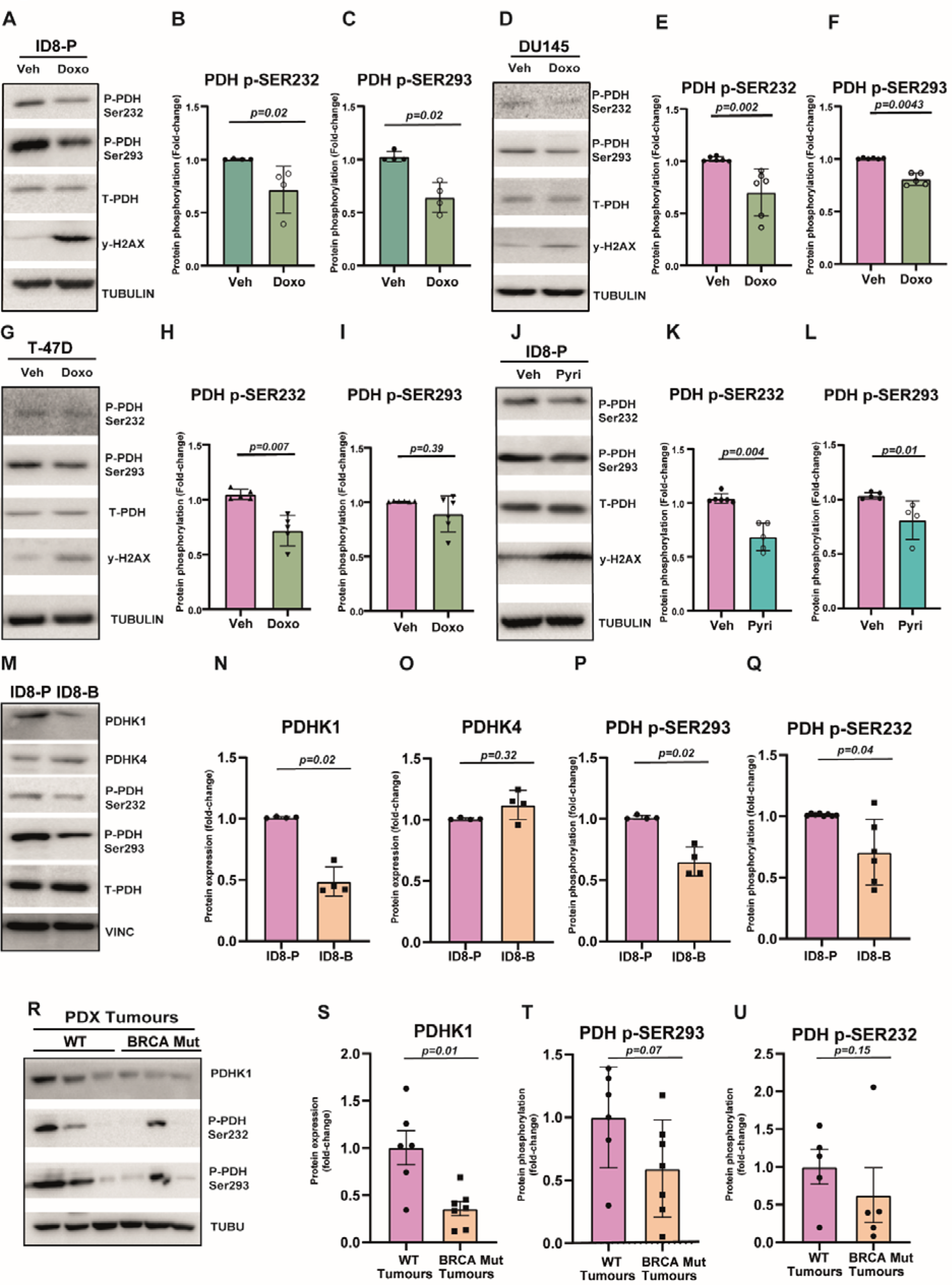
The accumulation of DSB increase the activity of the PDH promoting an increased OXPHOS metabolism by downregulating PDHK1. A) Western blot assessing the phosphorylation of the PDH on serine 232 and 293, total PDH (T-PDH), y-H2AX and tubulin (as a loading control) in ID8-*Trp53* KO cells (ID8-P), pretreated with vehicle (DMSO) or doxorubicin 2µM (Doxo) for 24 hours. A representative image is shown. **B- C)** Quantification of phospho-serine 232 and 293 in PDH shown in A). Tubulin was used as a loading control. **D)** Western blot assessing the phosphorylation of the PDH on serine 232 and 293, total PDH (T-PDH), y-H2AX and tubulin (as a loading control) in DU145 cells, pretreated with vehicle (DMSO) or doxorubicin 2µM (Doxo) for 24 hours. A representative image is shown. **E-F)** Quantification of phospho-serine 232 and 293 in PDH shown in D). Tubulin was used as a loading control. **G)** Western blot assessing the phosphorylation of the PDH on serine 232 and 293, total PDH (T-PDH), y-H2AX and tubulin (as a loading control) in T-47D cells, pretreated with vehicle (DMSO) or doxorubicin 2µM (Doxo) for 24 hours. A representative image is shown. **H-I)** Quantification of phospho-serine 232 and 293 in PDH shown in G). Tubulin was used as a loading control. **J)** Western blot assessing the serine 232 and 293 phosphorylation of the PDH, total PDH, y-H2AX and tubulin (as a loading control) in ID8-*Trp53* KO cells (ID8-P), pretreated with vehicle (Saline) or pyridostatin 10µM (Pyri) for 24 hours. A representative image is shown. **K-L)** Quantification of phospho-serine 232 and 293 in PDH shown in J). Tubulin was used as a loading control. **M)** Western blot assessing the expression of PDHK1 and 4, and the phosphorylation of the PDH in serine 232 and 293, total PDH and vinculin (loading control) in ID8-*Trp53* KO cells (ID8-P) and ID8-*Trp53 Brca2* KO cells (ID8-B). A representative image is shown. **N-Q)** Quantification of the different proteins and phosphorylation shown in M). Vinculin was used as a loading control. **R)** Western blot assessing the expression of PDHK1, the phosphorylation in serine 232 and 293 of PDH and tubulin (as a loading control) in HGSOC non mutated *BRCA*1/2 tumours (WT) or mutated in *BRCA1/2* (BRCA mut) growth as PDX in mice. A representative image is shown. **S-U)** Quantification of the different proteins and phosphorylation shown in R). Tubulin was used as a loading control. Data are presented Mean ± SD unless otherwise specified, and each graph dot represents an independent biological replicate. Significant differences were assessed using Mann- Whitney U-test when comparing between two groups and considered when P<0.05. The p value is indicated in each figure.

The phosphorylation of PDH emerged as a critical factor regulating the energetic metabolism upon DSB induction. Thus, we sought to identify the specific regulatory factors of the PDH axis that contribute to this regulation. We initially focused on chronic DNA damage in ovarian cancer by comparing ID8-P and ID8-B cells. We observed reduced mRNA expression of *Pdk1* and *Pdk4* genes in the BRCA model (Sup. Fig 3A), and PDHK1 was further confirmed by Western blot as well (Fig. 3M-O). Furthermore, consistent with the decrease on PDHK1, the phosphorylation of PDH at serine 293, which has the strongest correlation with its activity, and at serine 232, which is specific for the PDHK1, were both decreased (Fig 3M, P-Q). Similar patterns were observed in a SKOV3 *BRCA2* mutated model that we previously characterized^21^ (Sup. Fig. 3 B-F). Therefore, these evidences reinforce the importance of the PDH as a main regulator of the energetic metabolism towards maintaining a Warburg phenotype or shifting to an OXPHOS metabolism.

To confirm the implication of the PDHKs, we analysed the OCR/Glycolysis rate after treating ID8-P cells with a PanPDHK inhibitor (PanPDHKi) to increase PDH activity. This treatment led to an increased OCR/glycolysis ratio (Sup. Fig. 3G). Conversely, the use of a mitochondrial pyruvate carrier inhibitor (MPCi) in the ID8-B model, which impedes pyruvate entry into the mitochondria, resulted in a decreased OCR/glycolysis ratio (Sup. Fig. 3H). In addition, we examined the expression of PDHK isoforms by qPCR as well as PDHK1 and the phosphorylation of PDH by western blot in different PDX tumour samples from HGSOC patients. These samples were derived from patients with *P53* gene mutations (WT tumours) and patients with both *P53* and *BRCA1/2* gene mutations (BRCA Mut tumours). Interestingly, BRCA mutated tumours exhibited decreased *PDK1* mRNA (Sup. Fig. 3I-L) and PDHK1 protein and lower PDH phosphorylation (Fig. 3R-U). This provided further confirmation of the significance of the PDHK/PDH axis in regulating carbon flux and energy metabolism in ovarian cancer.

### HIF-1α downregulation after DSB damage is responsible for the metabolic shift towards OXPHOS

Next, our focus turned to identify the regulatory network controlling PDHK1 expression, as it is an enzyme under transcriptional control^24^. To this end, we conducted a comprehensive analysis of the *PDK1* promoter, seeking transcription factors with validated binding to specific promoter sites and known to be involved in the regulation of glycolysis using the ChIP-Atlas database^28^. Then, we correlated the expression levels of all the transcription factors identified with the HRD index using the Knijnenburg HRD score^27^(Sup. Fig. 4A-C) using publicly available mRNA data from the TGCA. This analysis unveiled two promising candidates: HIF-1 (Hypoxia-inducible factor 1) and BACH1 (BTB and CNC homology 1) that showed a remarkable link with the HRD scores. Consistently, the expression of a HIF1 signature was lower in samples with high HRD scores (Sup. Fig. 4D), suggesting a reduced expression and activity of this transcription factor in tumours with chronic HRD. To further confirm our bioinformatic findings, we examined the mRNA and protein of both transcription factors in our cell models. BACH1 and HIF1α expression was indeed decreased both at the mRNA and the protein level, in the BRCA mutated models (Fig. 4A-F; Sup. Fig. 4E-G). Furthermore, we sought to validate these results in ovarian tumours. For this, we assessed by qPCR in our PDX cohort the expression of both transcription factors and by IHC the protein expression in a retrospective series of high-grade serous epithelial ovarian carcinomas with or without mutations in *BRCA1* and *BRCA2* genes (Fig. 4G-L). Overall, our findings further supported the notion that these transcription factors could be involved in the regulation of the observed metabolic adaptations.

**Figure 4.**
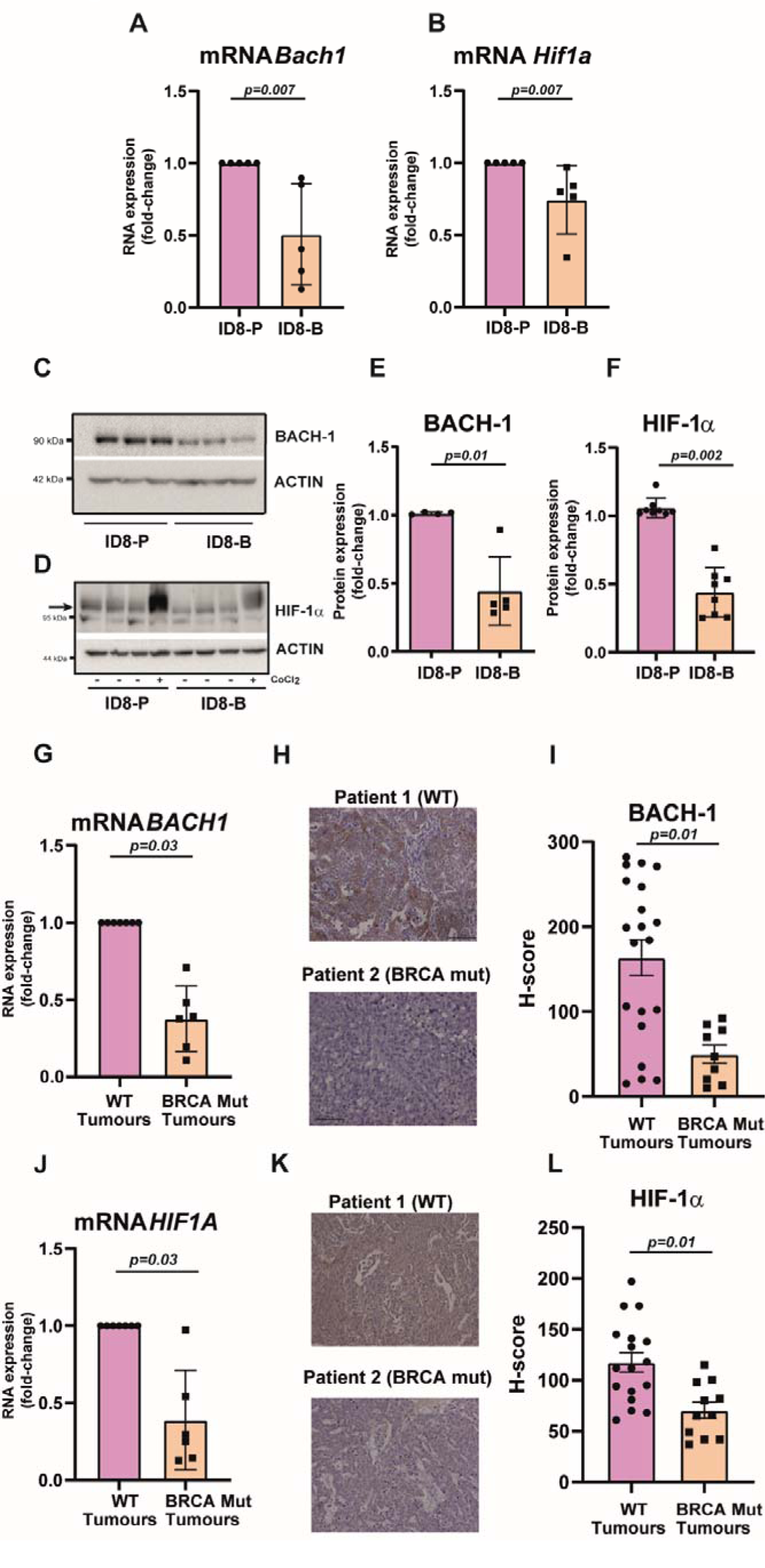
Glycolytic transcription factors are downregulated in HRD defective models A-B) mRNA levels of *Bach1* gene and *Hif1a* gene in ID8-*Trp53* KO cells (ID8-P) and ID8- *Trp53 Brca2* KO cells (ID8-B). *Actin* mRNA levels was used as control. **C)** Western blot assessing the expression of BACH-1 and β-actin (as a loading control) in ID8-*Trp53* KO cells (ID8-P) and ID8-*Trp53 Brca2* KO cells (ID8-B). A representative image is shown. **D)** Western blot assessing the expression of Hif-1α and β-actin (as a loading control) in ID8-*Trp53* KO cells (ID8-P) and ID8-*Trp53 Brca2* KO cells (ID8-B). Cells cultured with 0.2 mM of CoCl_2_ for 24h were used as a positive control of the presence of HIF-1α. A representative image is shown. **E-F)** Quantification of the different proteins shown in C and D). Actin was used as a loading control. **G)** mRNA levels of *BACH1* genes in HGSOC tumours. *Actin* mRNA levels were used as control (represented Mean ± SEM). **H)** IHC against BACH-1 in FFPE HGSOC human samples. Two representative images (one *BRCA* WT, one *BRCA1/2* mutated) are shown. **I)** Quantification (H-score) of the IHC shown in H) represented by the Mean ± SEM. **J)** mRNA levels of *HIF1A* gene in HGSOC tumours. Actin mRNA levels were used as control (represented Mean ± SEM). **K)** IHC against Hif- 1α in FFPE HGSOC human samples. Two representative images (one *BRCA* WT, one *BRCA* mutated) are shown. **L)** Quantification (H-score) of the IHC shown in K) represented Mean ± SEM. Data are presented Mean ± SD unless otherwise specified, and each graph dot represents an independent biological replicate. Significant differences were assessed using Mann-Whitney U-test when comparing between two groups and considered when P<0.05. The p value is indicated in each figure.

To elucidate the role of either transcription factor, firstly we initially knocked out the *Bach1* gene in the ID8-P model using Crispr/Cas9 technology. This intervention did not result in apparent modulation in PDHK1 protein levels or in PDH phosphorylation (Sup. Fig. 5A-D). In addition, the OCR/Glycolysis rate (Sup. Fig. 5E), was unchanged. This suggested that BACH-1 does not play a major role in the metabolic regulation of PDH in our model. Then, we silenced HIF-1α using shRNAs (Fig 5A) in the ID8-P background. Silencing was validated by measuring the expression of *Hif1a* and well-established HIF- 1 readouts (Sup Fig 5F). We observed a decrease in the expression of PDHK1 at mRNA and protein levels (Fig 5B, Sup Fig 5F) and in the phosphorylation of PDH serine 232 (Fig. 5A-D). Furthermore, the OCR/glycolysis ratio (Fig. 5E) was increased after *Hif1a* silencing, suggesting increased PDH activity as previously shown (Sup. Fig. 3G-H). We confirmed the implication of HIF-1α in the control of PDH activity using PX-478, a well-known HIF-1 inhibitor^36^, which caused a drop in PDHK1 expression and in the phosphorylation levels of PDH Ser232 (Sup. Fig. 5G-I) and an increased OCR/glycolysis ratio (Sup. Fig. 5J). This led us to confirm our hypothesis that HIF-1 is the main regulator of PDH activity in this context. Finally, to query if in the acute induction of DSBs impacts on HIF-1 expression on non-HR deficient models we validated if HIF-1α was downregulation upon DSB induction. Results observed were consistent with acute DNA damage, either by inhibiting topoisomerases (Fig 5F-H, Sup. Fig. 6A-F) or by the stabilization of G-quadruplex cells (Sup. Fig. 6G-I), also controlling metabolic plasticity via a common molecular mechanism through the HIF-1-PDHK1-PDH axis. Moreover, we observed the same downregulation of HIF-1 upon DSBs in cell lines from prostate (Fig 6I-K) and breast cancer (Fig 6L-N). Therefore, our results indicated that HIF-1 is a key regulator of metabolic plasticity in our model.

**Figure 5.**
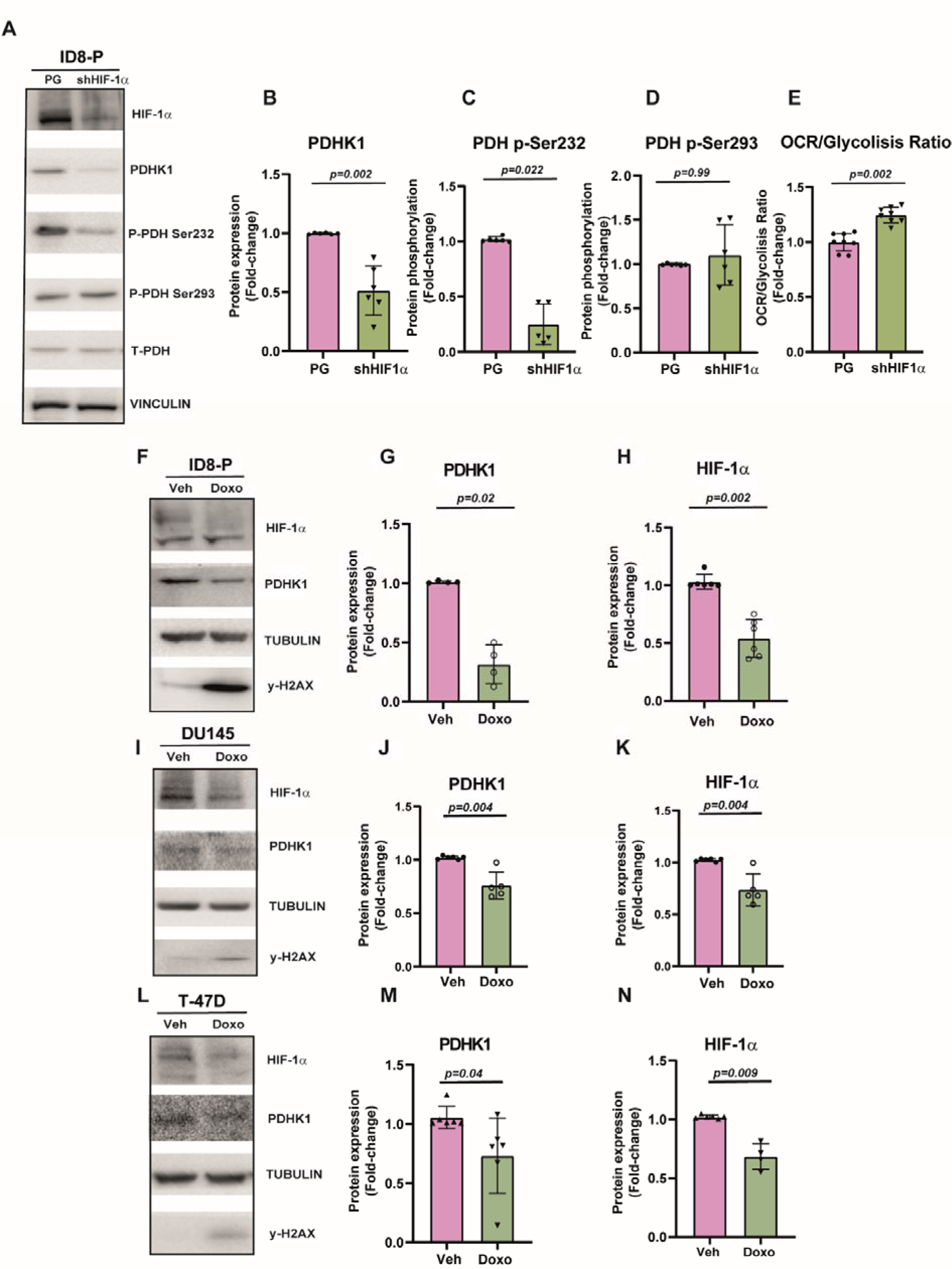
Downregulation of HIF-1. α **is needed to increase the oxidative metabolism. A)** Western blot assessing the expression of PDHK1, the phosphorylation of serine 232 and 293 of PDH, total PDH and vinculin (as a loading control) in control (ID8 P53-GFP, PG) and *Hif1a* knockdown cells (shHIF1α). A representative image is shown. **B-D)** Quantification of the different proteins shown in A). Vinculin was used as a loading control. **E)** OCR/Glycolysis ratio is shown in control (ID8 P53-GFP, PG) and *Hif1a* knockdown cells (shHIF1α). **F)** Western blot assessing the expression of HIF-1α, y-H2AX and tubulin (as a loading control) in ID8-*Trp53* KO cells (ID8-P), pretreated with vehicle (DMSO) or doxorubicin 2µM (Doxo) for 24 hours. A representative image is shown. **G- H)** Quantification of phospho-serine 232 and 293 in PDH shown in F). Tubulin was used as a loading control. **I)** Western blot assessing the expression of PDHK, HIF-1α, and the phosphorylation of H2AX in DU145 cells, pretreated with vehicle (DMSO) or doxorubicin 2µM (Doxo) for 24 hours. A representative image is shown. **J-K)** Quantification of the different proteins shown in I). Tubulin was used as a loading control. **L)** Western blot assessing the expression of PDHK, HIF-1α, and the phosphorylation of H2AX in DU145 cells, pretreated with vehicle (DMSO) or doxorubicin 2µM (Doxo) for 24 hours. A representative image is shown. **M-N)** Quantification of the different proteins shown in L). Tubulin was used as a loading control. Data are presented Mean ± SD unless otherwise specified, and each graph dot represents an independent biological replicate. Significant differences were assessed using Mann-Whitney U-test when comparing between two groups and considered when P<0.05. The p value is indicated in each figure.

**Figure 6:**
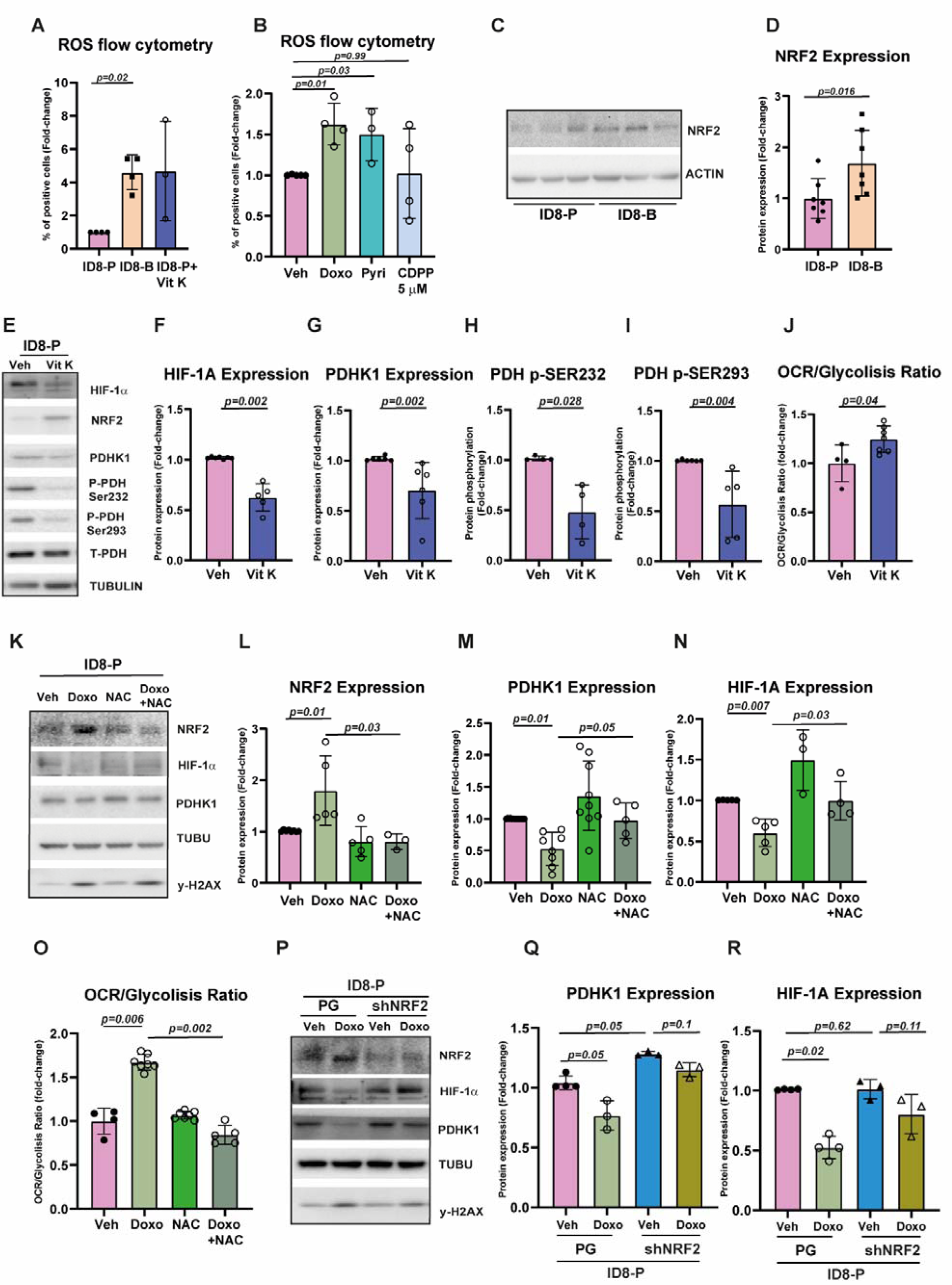
ROS induced by DSB activate NRF2 in order to downregulate the HIF- 1/PDHK1 axis. A) Quantification of cytosolic ROS in ID8-*Trp53* KO cells (ID8-P), ID8- *Trp53 Brca2* KO cells (ID8-B) and ID8-P cells pretreated with Vitamin K3 10µM (Vit K), as a positive control. **B)** Quantification of cytosolic ROS in ID8-*Trp53* KO cells pretreated with vehicle (DMSO), Doxorubicin 2µM (Doxo), pyridostatin 10µM (Pyri) or cisplatin 5µM (CDPP) for 24 hours. **C)** Western blot assessing the expression of NRF2 and β-actin (as a loading control) in ID8-*Trp53* KO cells (ID8-P) and ID8-*Trp53 Brca2* KO cells (ID8-B). A representative image is shown. **D)** Quantification of NRF2 shown in C). Tubulin was used as a loading control. **E)** Western blot assessing the expression of HIF- 1α, NRF2, PDHK1, the phosphorylation of serine 232 and 293 of PDH, total PDH and Tubulin (as a loading control) in ID8-*Trp53* KO cells (ID8-P), pretreated with vehicle (DMSO) or Vitamin K3 10µM (Vit K) for 24 hours. A representative image is shown. **F-I)** Quantification of the different proteins shown in E). Tubulin was used as a loading control. **J)** OCR/Glycolysis ratio is shown. ID8-*Trp53* KO cells (ID8-P) were previously treated 24 hours with vehicle (DMSO) or Vitamin K3 10µM (Vit K). **K)** Western blot assessing the expression of NRF2, PDHK1, HIF-1α, y-H2AX and Tubulin (as a loading control) in ID8-*Trp53* KO cells (ID8-P), pretreated with vehicle (DMSO), Doxorubicin 2µM (Doxo), N-acetyl cysteine 0.5mM (NAC) or the combination of both for 24 hours. A representative image is shown. L-N) Quantification of the different proteins shown in K). Tubulin was used as a loading control. **O)** OCR/Glycolysis ratio is shown. ID8-*Trp53* KO cells (ID8-P) were previously treated 24 hours with vehicle (DMSO), Doxorubicin 2µM (Doxo), N-acetyl cysteine 0.5mM (NAC) or the combination of both. **P)** Western blot assessing the expression of PDHK1, HIF-1α, NRF2 and tubulin (as a loading control) in control (ID8 P53-tdTomato, PG) and *Nfe2l2* knockdown cells (shNRF2). A representative image is shown. **Q-R)** Quantification of the different proteins shown in P). Tubulin was used as a loading control. Data are presented Mean ± SD unless otherwise specified, and each graph dot represents an independent biological replicate. Significant differences were assessed using Mann-Whitney U-test when comparing between two groups and considered when P<0.05. The p value is indicated in each figure.

### DSB-induced ROS activate NRF2 and promote the downregulation of the HIF- 1/PDHK1 axis

Next, we addressed the mechanistic link between DSB accretion and HIF-1 regulation. We initially hypothesized that DSB response pathways might be implicated in the regulation of HIF-1 observed. However, the inhibition of canonical DSB response pathways did not limit the degradation of HIF-1 observed (Sup. Fig. 7A-B). Alternatively, for example, DSB can indirectly stimulate the production of reactive oxygen species (ROS) in the cell^37–39^, hence we focused on DSB mediated ROS signalling as a transduction mechanism. Indeed, cytosolic ROS increased in parallel with DSB accumulation in our cellular models. For example, ID8-B cells had higher levels of ROS than ID8-P cells, suggesting that cells with chronic DNA damage have more oxidative stress (Fig 6A). Direct effects of DSB inducers on ROS levels were also apparent when treating ID8-P cells with pyridostatin or doxorubicin (Fig 6B), whereas non-DSB inducing DNA damaging agents such as CDPP 5μM, did not increase ROS (Fig 6B), further suggesting that this signal is specific for a DSB stimulus either chronically or acutely accrued. Consistently, we observed that our cell lines with HRD presented higher levels of NRF2, a ROS sensor (Fig 6C-D, Sup. Fig 7C-D). Finally, ROS production in ID8-P cells using Vitamin K3 (Fig 6E-I) was sufficient to produce the metabolic and molecular shifts that we had observed with the treatment with DSB inducers, namely downregulation of the HIF-1/PDHK1 axis (Fig 6F-I) and increasing OXPHOS metabolism (Fig 6J).

As NRF2 was clearly stimulated in this condition, we hypothesize that NRF2 might be mediating the molecular events associated with DSB cellular metabolism adaptations. Further validating this hypothesis, we measured increased levels of NRF2 after treatment with DSB inducers (Fig 6K-L, Sup. Fig. 7E-H), and this induction was reverted with incubation with NAC, an antioxidant (Fig 6K-N). To note, the combination of doxorubicin with NAC, the downregulation of HIF-1/PDHK1 axis was prevented (Fig 6M-N) and the metabolic shift towards oxidative metabolism was also abrogated (Fig 6O). To further demonstrate that this effect was NRF2-dependent, we use DMF (Sup Fig 7I-N), a NRF2 stabilisation agent^40^, that does not produce ROS (Sup Fig 7O). We, again, observed a similar increase in OXPHOS metabolism accompanied by the downregulation of HIF-1/PDHK1 axis (Sup Fig 7I-N). To fully confirm the implication of NRF2, we silenced its expression in ID8-P and ID8-B cells. First, we observed that upon doxorubicin treatment, ID8-P cells expressing the shRNA against NRF2 could not promote the expression of NRF2 readouts that were upregulated upon doxorubicin treatment in control cells (Sup Fig 7P). Second, the HIF-1/PDHK1 axis did not respond to doxorubicin in the absence of NRF2 (Sup Fig 7P and Fig. 6P-R), and thirdly, ID8-B cells expressing the shRNA against NRF2, presented a rebound expression of HIF-1 and PDHK1 (Sup. Fig. 7Q-S), suggesting that the chronic DNA damage effects on this pathway were partially reverted, and demonstrating that NRF2 is a key mediator in this metabolic shift.

**Figure 7.**
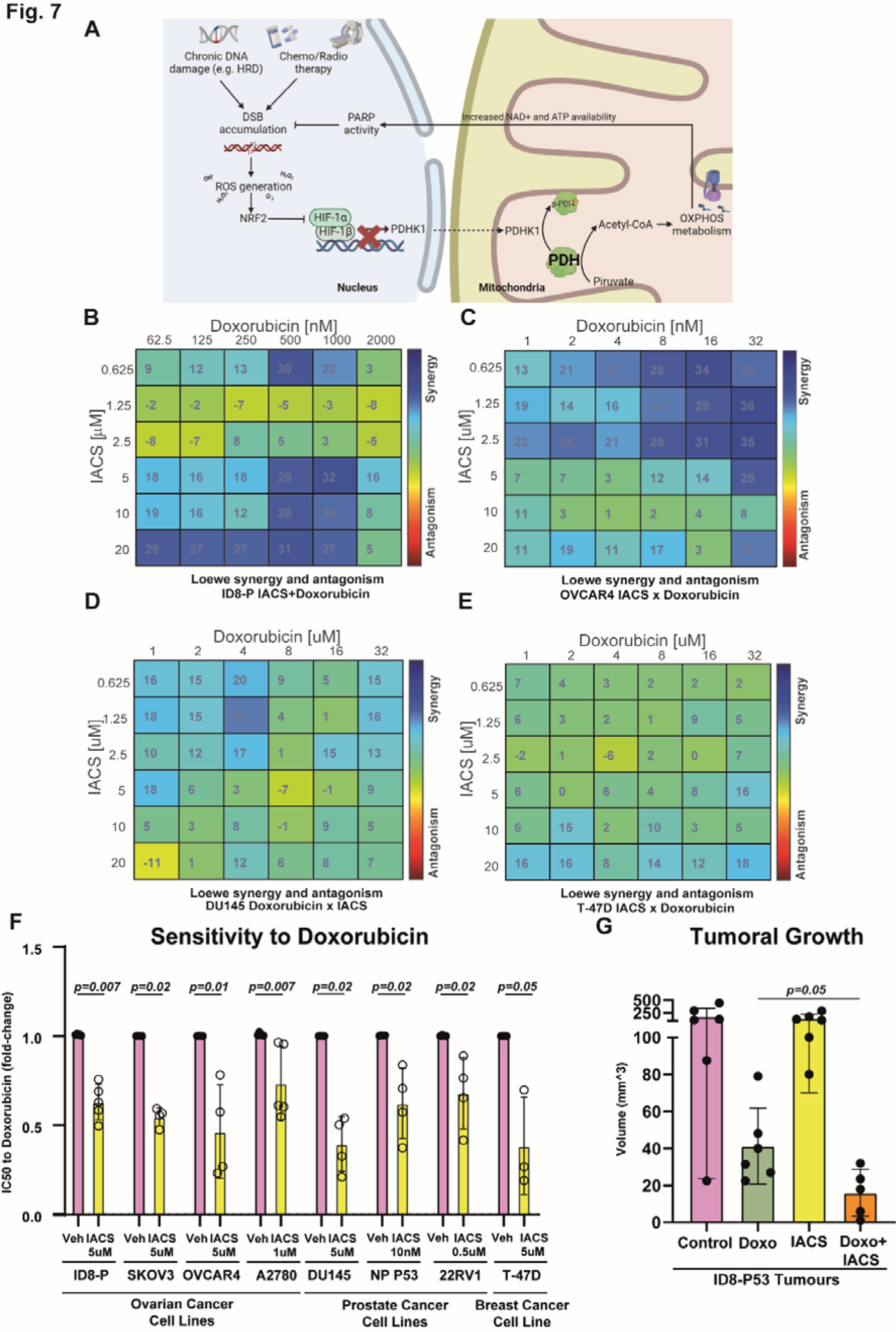
Doxorubicin and OXPHOS inhibitors have a synergistic effect in vitro shared among several cancer cell types and maintains their efficacy in vivo. A) Scheme of the process described in the present work created with BioRender.com **B-E)** Synergy matrix of ID8-*Trp53* KO cells (ID8-P) (B), OVCAR4 (C), DU145 (D), T-47D (E) incubated with different doses of doxorubicin and IACS-10759. **F)** Graph showing the IC50 to doxorubicin. ID8-*Trp53* KO cells (ID8-P), SKOV3N (SKOV3), OVCAR4, A2780, DU145, NP P53, 22Rv1 and T-47D cells were treated with vehicle (DMSO) or IACS-10759 (5µM for ID8-P, SKOV3, OVCAR4, DU145 and T-47D; 1µM for A2780; 0.5µM for 22Rv1; and 10nM for NP P53) and then with increasing doses of doxorubicin for 72 hours. IC50 was assessed with the non-linear regression tool from GraphPad Prism 8. **G)** Graph assessing the tumoral volume of the primary tumour. Mice were treated with control (Control; 100 µL of saline 5% glucose once a week with a intraperitoneal injection, 200µL of carboxymethyl cellulose 0.5% in a 5 days treated-2 day rest regime by oral gavage) doxorubicin (Doxo, 2mg/kg of Caelyx [liposomal pegylated doxorubicin] dissolved in saline 5% glucose once a week with a intraperitoneal injection, 200µL of carboxymethyl cellulose 0.5% in a 5 days treated-2 day rest regime by oral gavage), IACS-10759 (IACS; 100 µL of saline 5% glucose once a week with a intraperitoneal injection, 0.3mg/kg of IACS-10759 dissolved in carboxymethyl cellulose 0.5% in a 5 days treated-2 day rest regime by oral gavage) and doxorubicin+IACS-10759 (Doxo+IACS; 2mg/kg of Caelyx [liposomal pegylated doxorubicin] dissolved in saline 5% glucose once a week with a intraperitoneal injection 0.3mg/kg of IACS-10759 dissolved in carboxymethyl cellulose 0.5% in a 5 days treated-2 day rest regime by oral gavage). Data are presented Mean ± SD unless otherwise specified, and each graph dot represents an independent biological replicate. Significant differences were assessed using Mann-Whitney U-test when comparing between two groups and considered when P<0.05. The p value is indicated in each figure.

### The combination of OXPHOS inhibitors with DSB inducers has a synergistic effect on tumour growth inhibition in vitro and in vivo

In view of our intriguing results, which indicate that NRF2/HIF-1 are a key rheostat in the control of the metabolic plasticity (Fig. 7A) in our models and since the use of DNA damage inducers stands as one of the main strategies for tumour cell death promotion, we hypothesized that combined chemotherapy with NRF2 activators or HIF-1 inhibitors could hold therapeutic benefits. To validate this hypothesis, we conducted synergy assays to assess the effectiveness of these combinations. In the synergy analysis, we generated a matrix with the viability of each combination compared to the control condition (Sup. Fig. 8A). Subsequently, we used the Combenefit software to calculate the synergy effect of each combination, as previously described^31^. For this analysis, doxorubicin was used, which is administered to platinum resistant ovarian cancers and metastatic breast cancers among several other cancer types. Regarding HIF-1 inhibition PX-478 was used, and for NRF2 activation cells were incubated with DMF. The assay showed that the combination of both inhibitors with doxorubicin had a summative effect but was not synergistic (Sup. Fig. 8B-C). This was also confirmed by performing the IC50 to doxorubicin (Sup. Fig. 8D-E) upon the presence of the HIF-1 inhibitor or the NRF2 activator, were we observed a mild reduction. Albeit unappreciated initially, we reason that having both inhibitors promoting a similar effect reducing HIF-1 activity, could not yield a synergistic activity.

Therefore, we designed a strategy based on targeting new vulnerabilities triggered by DSB-inducing treatments. Hence, as DSB promote an increased dependency on OXPHOS metabolism, we hypothesized that the combination of DSB induction with OXPHOS inhibitors could be synergistic. For OXPHOS inhibition we used IACS-10759, a last generation inhibitor of the mitochondrial respiratory chain complex I. Remarkably, the assay exhibited high levels of synergy (Fig. 7B). To validate and reinforce these findings, we performed the doxorubicin+IACS synergy assay in a different ovarian cancer cell line, OVCAR4 (Fig 7C) and in a prostate (Fig 7D) and breast cancer (Fig 7E) cell lines. Furthermore, we conducted additional validations by measuring the IC50 of doxorubicin in a panel of different cell lines (Fig. 7F). For all cell lines tested, we observed increased sensitivity to doxorubicin in the presence of IACS regardless of the HR status of the cell line (Sup. Table 2). To validate the effectiveness of the combined treatment in an *in vivo* setting, we treated mice with ID8-P with a combination of chemotherapy (doxorubicin) and OXPHOS inhibition (IACS) (Fig. 7G). Orthotopic implantation of 2 million ID8-P53 cells in the right ovary of immunocompetent C57BL6J mice was performed, and after 4 weeks of tumour growth, we initiated the treatment. Given the potential secondary effects of IACS and its limited efficacy as a monotherapy, we opted for a very low dose (0.3 mg/kg) that was 8 to 30 times lower than those used in previous studies^41–43^. We hypothesized that a reduced dosage might still impair tumour growth while avoiding any significant secondary effects when combined with doxorubicin. For chemotherapy, we used pegylated liposomal doxorubicin (Caelyx, 2mg/kg), the one commonly used in ovarian cancer clinics. Throughout the course of the experiment, we closely monitored the mice’s weight, glucose, and lactate blood levels to identify adverse effects related to the inhibition of hepatic gluconeogenesis and skeletal muscle function secondary to OXPHOS inhibition, but we did not detect any significant changes attributable to the treatment (Sup. Fig. 9A-C), confirming the absence of secondary effects in the mice. Also, as anticipated, low doses of IACS monotherapy did not produce any significant effect on tumoral growth. In contrast, doxorubicin alone reduced tumoral growth by 80%. However, the combination of doxorubicin+IACS had a dramatic effect, reducing total tumoral volume by 95%, which was a remarkable improvement compared to the doxorubicin monotherapy group (Fig. 7G).

In conclusion, these results offer a promising therapeutic strategy for treating different cancer types. Importantly, the reduced dosage of IACS used in the combination regimen minimized potential side effects without compromising treatment efficacy, making it an attractive option for further exploration in clinical settings.

## DISCUSSION

Metabolic plasticity, defined as the ability of cells to utilize the same substrate in different ways based on their specific needs, plays a crucial role in cancer cell survival^44^. In our current study, we have demonstrated that when tumours encounter DSB in their genome, caused either by chronic DNA damage (e.g., deficiency in *BRCA1/2* or related repair genes) or by treatment with conventional anti-tumoral agents (e.g., doxorubicin, irinotecan, or radiotherapy), their metabolic plasticity facilitates a shift in carbon and energy utilization. Specifically, tumours transit from a partial but rapid oxidation of glucose, known as the Warburg phenotype, to its complete mitochondrial oxidation. This change to complete mitochondrial oxidation is highly efficient, allowing tumour cells to produce elevated levels of ATP and restore NAD^+^ levels, both essential for sustaining DNA repair through the A-EJ pathway. Interestingly, this increase in glucose consumption has been described by our group and others as a generalized adaptive response, at least in the initial phase, to various stressors in the absence of DSBs to establish a favourable energetic environment for survival^45^.

However, DSB induction not only boosts glucose consumption but also enhances its complete oxidation. This unique metabolic shift is further enabled by a reduction in proliferation. Unlike normal cancer cells which predominantly follow a Warburg metabolism to generate building blocks (e.g. nucleotides) for a fast proliferation, cells with damaged DNA redirect their resources to counteract the redox an energetic imbalance caused by DNA repairing mechanisms thus showing an attenuated proliferation^46^. Importantly, our findings reveal that this adaptive response is a general mechanism observed in different cancer types. Previous studies in cell lines have also observed an increase in OXPHOS consumption when DNA damage is induced^47,48^, indicating the critical role of OXPHOS metabolism as part of the DNA damage response mechanisms^49–51^.

The metabolic pivot induced by DSB, leading to an increase in OXPHOS metabolism, is mediated by the accumulation of cytoplasmatic ROS that induce the activation of NRF2 and a reduction in HIF-1α levels, as observed at both the RNA and protein levels. While BACH-1 also decreases in response to DSB, our findings suggest that this transcription factor may play a less critical and potentially redundant role in the process, although it has extensively been described a crosstalk with NRF2^52–54^. Interestingly, the decrease in HIF-1α or BACH-1 has been implicated in inducing OXPHOS in several other situations^55–57^ although the mechanism behind the coupling of these pathways are yet to be identified. Consistently, existing evidence suggests that the induction of the NRF2 transcription factor by ROS has been linked to the decrease in glycolytic metabolism and the increase in OXPHOS observed in other contexts^58^. Despite we cannot rule out the implication of other mechanisms, it is clear that maintaining low HIF-1 activity is essential to trigger the subsequent decrease in PDHK1, resulting in a drop in PDH phosphorylation, an increase in its activity, and ultimately leading to the complete mitochondrial oxidation of pyruvate. This metabolic shift appears to be a crucial adaptation to efficiently respond to DSB and maintain cellular function and survival.

The metabolic shift involving the upregulation of OXPHOS metabolism by increasing PDH activity and respiratory chain activity, particularly in HRD tumours, represent a previously unappreciated vulnerability that opens a promising therapeutic opportunity in these cancer subtypes. Our previous study demonstrated the effectiveness of metformin in treating HR-deficient tumours, while it showed limited efficacy in non-HRD tumours^21^. However, the clinical use of OXPHOS inhibitors as a monotherapy has faced challenges due to their high toxicity^42^. Therefore, their application should be restricted to combinations with other drugs targeting alternative pathways or radiotherapy, ensuring low non-toxic doses while maintaining high efficacy. In this regard, our current findings suggest that OXPHOS inhibitors (used 8 to 30 times lower than others^41–43^) can be used not only in HR-deficient tumours but also in non-HR-deficient tumours when combined with chemotherapy regimens or radiotherapy that induce DSB, such as the combination of doxorubicin with IACS-10759 presented in this work. This finding opens the possibility of using this combination not only in cases where standard treatments fail, such as using doxorubicin alone in platinum-resistant ovarian cancers, but also in situations where patients may have chemotherapy intolerance. In such cases, the use of low doses of the chemotherapeutic agent combined with agents like IACS-10759 could enhance treatment efficacy by achieving increased tumour reduction.

In summary, the present work sheds light on the widespread significance of OXPHOS metabolism in the DNA damage response, providing valuable insights into understanding the relationship between these two hallmarks of cancer. The ability to modulate the metabolism in response to DSB offers novel treatment options paving the way for potentially revolutionizing ovarian cancer treatment and may have broader implications for other cancer as well.

## ACKNOWLEDGEMENTS

This study has been funded by the Ministerio de Ciencia, Innovación y Universidades, which is part of the Agencia Estatal de Investigación (AEI), through the project SAF2017-85869-R and PID2020-117815RB-I00 (cofounded by the European Regional Development Fund (ERDF), a way to build Europe) to FV and BFU2015-66030-R and PID2019-106640RB-I00 to JCP; by the support of the Secretariat d’Universitats i Recerca of the Generalitat de Catalunya (2021 SGR 00184) to FV. FMJ was awarded with an FI-SDUR fellowship of the Agència de Gestió d’Ajuts Universitaris i de Recerca (AGAUR) which is part of the Departament de Recerca i Universitats of the Generalitat de Catalunya.

We particularly wish to acknowledge the collaboration of the patients. FMJ and FVC thank Dr. Alvaro Aytés (ProCure), Dr. Cristina Muñoz (Oncobell), and the rest of the Molecular Signalling Group (ProCure) for helpful discussions and suggestions. We thank the IDIBELL Scientific and Technical Services for the assessment in technical issues.

## AUTOR CONTRIBUTIONS

Conception and design: FM-J, FV, JCP. Acquisition of data (provided animals, acquired and managed patients, provided essential reagents, etc.): FM-J, AF, AL, PG, EB, AV, AB- P, MR. Analysis and interpretation of data (e.g., statistical analysis, biostatistics, computational analysis): FM-J, RE, MAP, MAP, JCP, FV. Writing, review, and/or revision of the manuscript: FM-J, EB, JCP, FV. Study supervision: JCP, FV

---

**Supplementary Figure 1:**
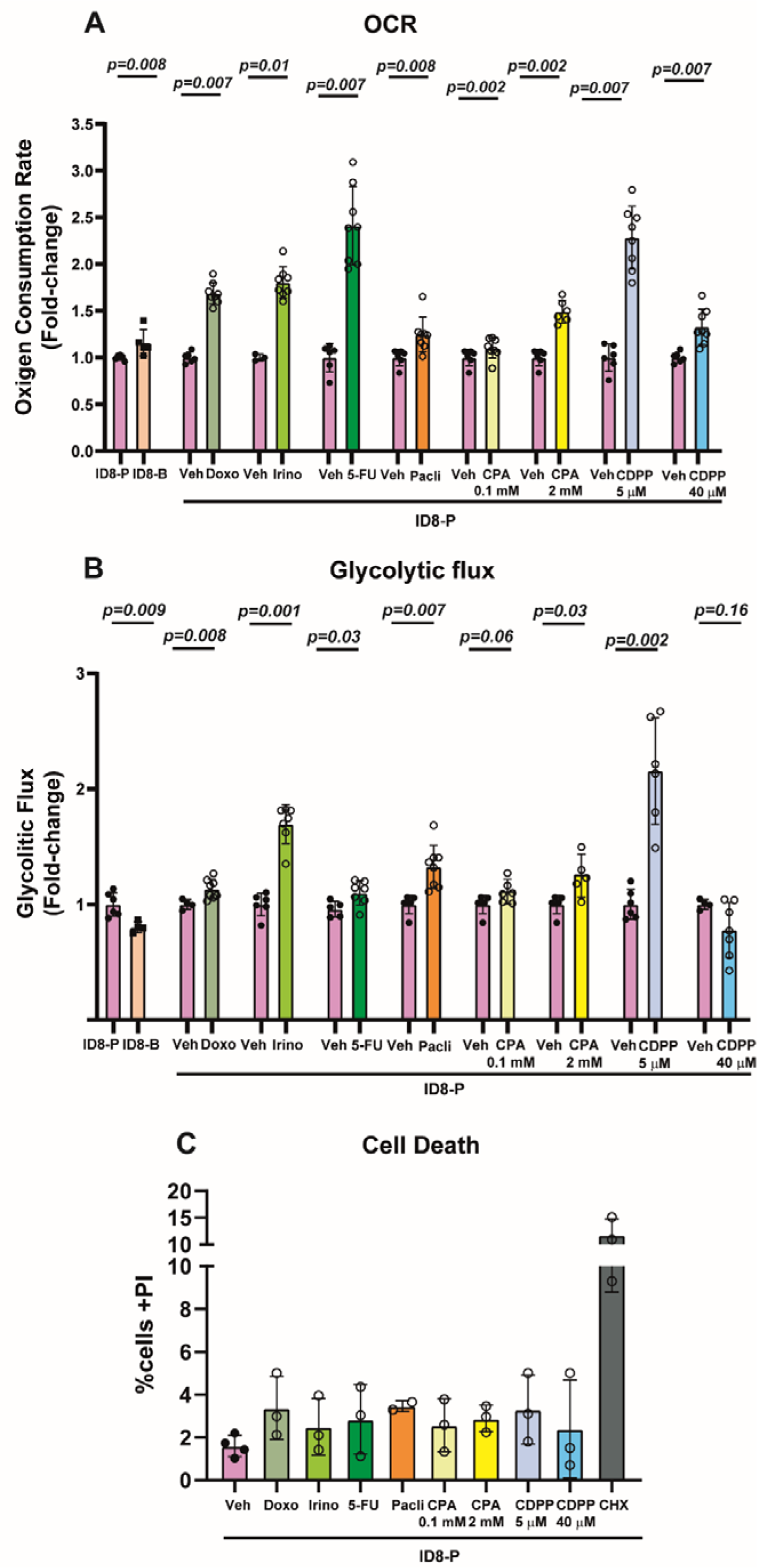
Chemotherapy increases the glycolytic flux and reduces cell number without affecting cell death. A) OCR is shown in ID8-*Trp53* KO cells (ID8-P) and ID8-*Trp53 Brca2* KO cells (ID8-B) and ID8-P cells pretreated for 24h with vehicle (DMSO), doxorubicin 2µM (Doxo), irinotecan 50µM (Irino), 5-fluorouracil 5µM (5-FU), paclitaxel 20nM (Pacli), ciclophosphamide 0.5mM or 2mM (CPA) or cisplatin 5µM or 40µM (CDPP). **B)** Glycolysis rate is shown in ID8-*Trp53* KO cells (ID8-P) and ID8-*Trp53 Brca2* KO cells (ID8-B) and ID8-P cells pretreated for 24h with vehicle (DMSO), doxorubicin 2µM (Doxo), irinotecan 50µM (Irino), 5-fluorouracil 5µM (5-FU), paclitaxel 20nM (Pacli), ciclophosphamide 0.5mM or 2mM (CPA) or cisplatin 5µM or 40µM (CDPP). **C)** Cell death measured as percentage of cells positive for propidium iodide is shown in ID8-*Trp53* KO cells (ID8-P) pretreated for 24h with vehicle (DMSO), doxorubicin 2µM (Doxo), irinotecan 50µM (Irino), 5-fluorouracil 5µM (5-FU), paclitaxel 20nM (Pacli), ciclophosphamide 0.5mM or 2mM (CPA), cisplatin 5µM or 40µM (CDPP) or 40µg/mL of cycloheximide (CHX) as a positive control. Data are presented Mean ± SD unless otherwise specified, and each graph dot represents an independent biological replicate. Significant differences were assessed using Mann-Whitney U-test when comparing between two groups and considered when P<0.05. The p value is indicated in each figure.

**Supplementary Figure 2:**
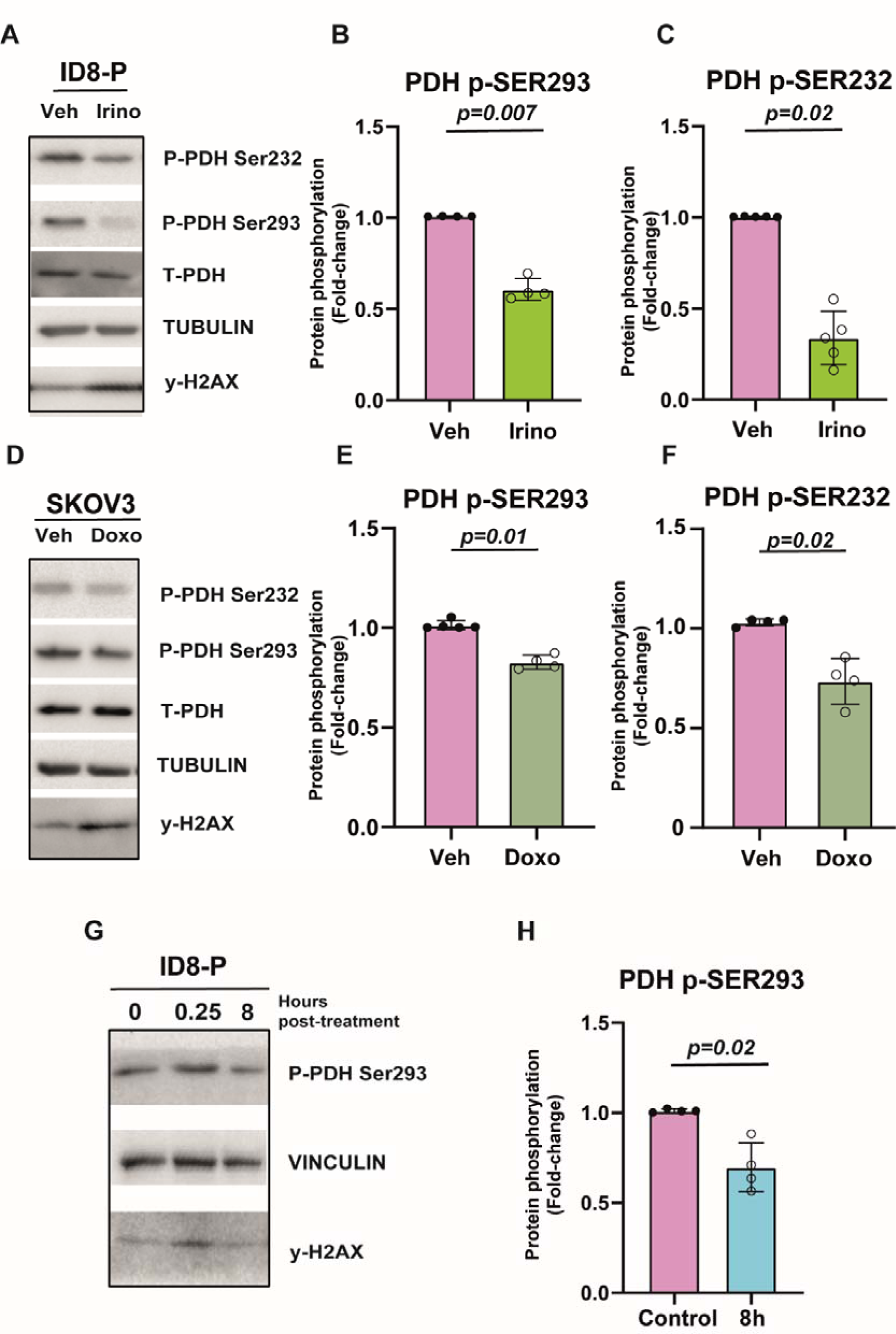
DBS induction promotes PDH activation. A) Western blot assessing the phosphorylation of serines 232 and 293 of the PDH, total PDH, γ-H2AX and tubulin (as a loading control) in ID8-*Trp53* KO cells (ID8-P), pretreated with vehicle (DMSO) or Irinotecan 50µM (Irino) for 24 hours. A representative image is shown. **B-C)** Quantification of the different proteins shown in A). Tubulin was used as a loading control. **D)** Western blot assessing the phosphorylation of serines 232 and 293 of the PDH, total PDH, γ-H2AX and tubulin (as a loading control) in SKOV3N cells, pretreated with vehicle (DMSO) or doxorubicin 2µM (Doxo) for 24 hours. A representative image is shown. **E-F)** Quantification of the different proteins shown in D). Tubulin was used as a loading control. **G)** Western blot assessing the phosphorylation of serine 293 of the PDH, γ-H2AX and vinculin (as a loading control) in ID8-*Trp53* KO cells (ID8-P), treated with a single dose of 6Gy of radiotherapy and collected at 0, 0.25 or 8 hours after the exposure. A representative image is shown. **H)** Quantification of serine 293 of the PDH is shown in G). Tubulin was used as a loading control. In each graph is represented by the Mean ± SD unless otherwise specified, and each dot represents an independent biological replicate. Significant differences were assessed using Mann-Whitney U-test when comparing between two groups and considered when P<0.05. The p value is indicated in every figure.

**Supplementary Figure 3.**
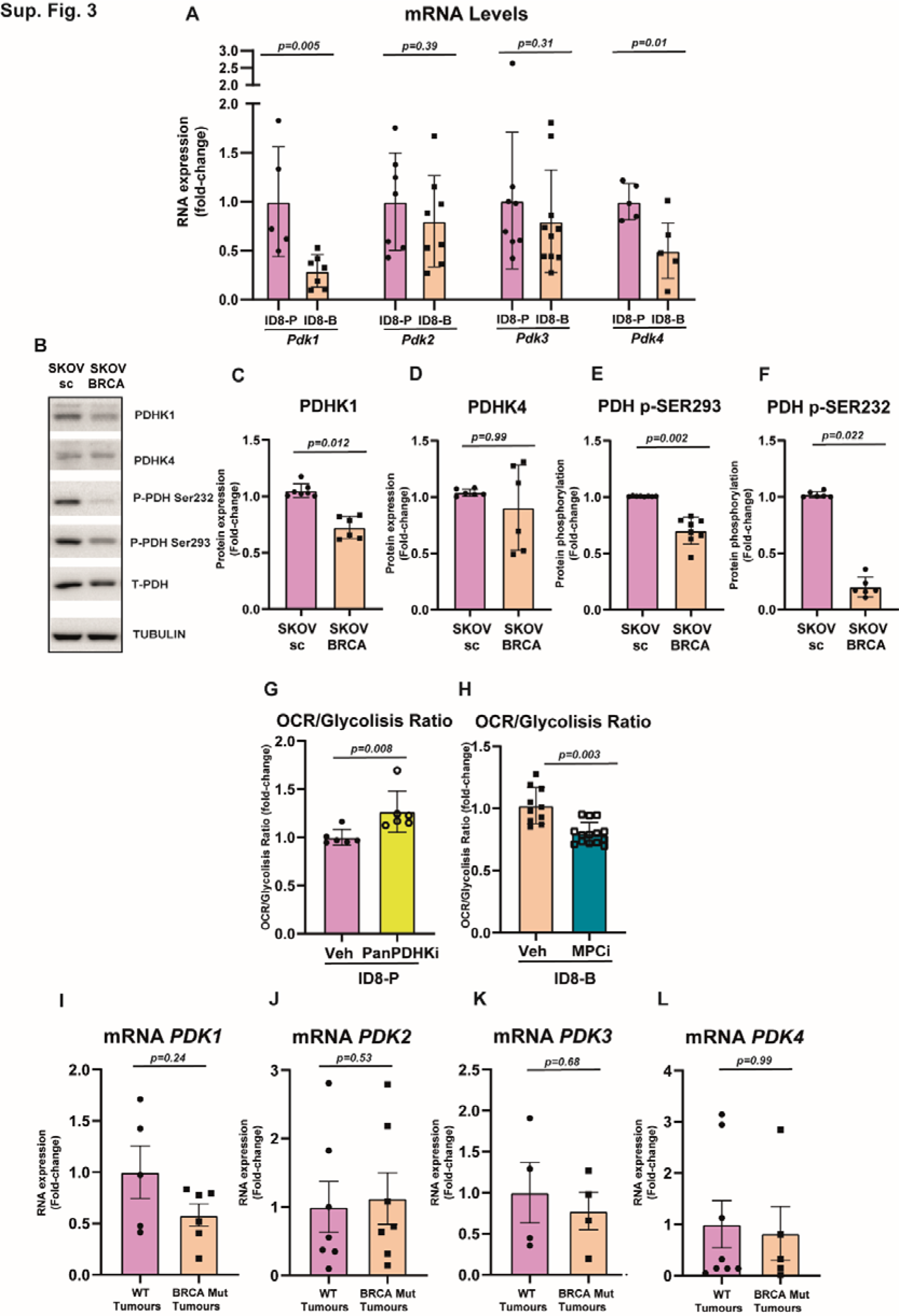
PDHK1/PDH axis regulation in other HR-deficient models. A) mRNA levels of each *Pdk* gene in ID8-*Trp53* KO cells (ID8-P) and ID8-*Trp53 Brca2* KO cells (ID8-B). *Actin* mRNA levels was used as control. **B)** Western blot assessing the expression of PDHK1 and 4, the phosphorylation in serines 232 and 293 of the PDH, total PDH and tubulin (as a loading control) in control SKOV3N cells (SKOV-sc) and SKOV3 *BRCA2* KO cells (SKOV-BRCA). A representative image is shown. **C-F)** Quantification of the different proteins and phosphorylation shown in B). Tubulin was used as a loading control. **G)** OCR/Glycolysis ratio is shown. ID8-*Trp53* KO cells (ID8-P) were previously treated 24 hours with VER-246608 500nM (PanPDHKi). **H)** OCR/Glycolysis ratio is shown. ID8-*Trp53 Brca2* KO cells (ID8-B) cells were previously treated 24 hours with GW604714 250nM (MPCi). **I-L)** mRNA levels of each *PDK* gene in HGSOC non mutated *BRCA*1/2 tumours (WT) or mutated in *BRCA1/2* (BRCA mut) growth as PDX in mice. *ACTIN* mRNA levels were used as control (represented by the Mean ± SEM). Data are presented Mean ± SD unless otherwise specified, and each graph dot represents an independent biological replicate. Significant differences were assessed using Mann-Whitney U-test when comparing between two groups and considered when P<0.05. The p value is indicated in each figure.

**Supplementary Figure 4.**
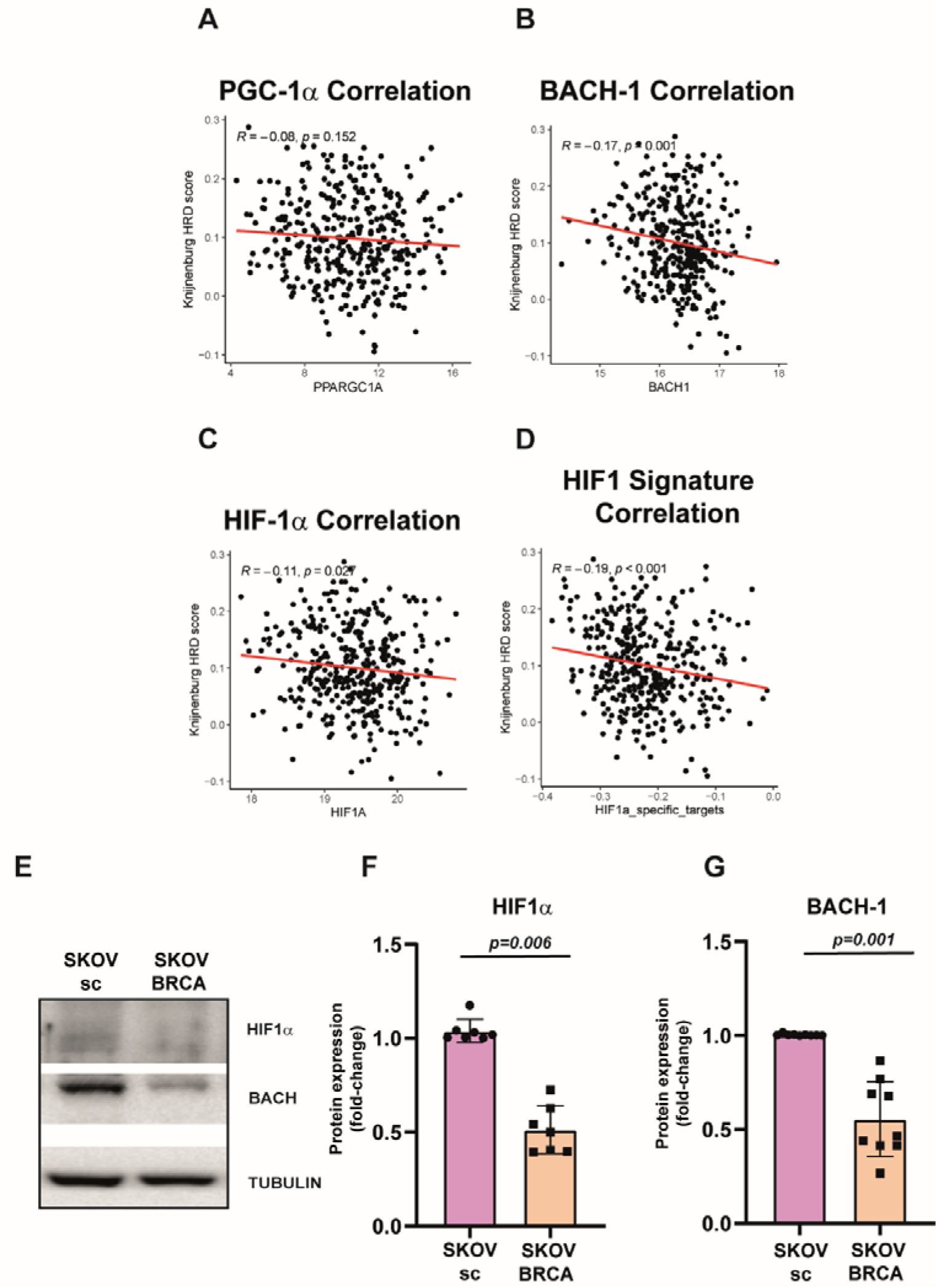
HIF-1. α **and BACH-1 expression in other HR-deficient models. A-C)** Bioinformatical analysis showing the mRNA expression of PGC-1α(A), *BACH1* (B), and *HIF1A* (C) genes correlated with the Knijnenburg HRD score. **D)** Bioinformatical analysis showing the correlation between the Knijnenburg HRD score and the score for the HIF-1α signature of each sample. **E)** Western blot assessing the expression of HIF-1α, BACH-1 and tubulin (as a loading control) in control SKOV3N cells (SKOV-sc) and SKOV3 *BRCA2* KO cells (SKOV-BRCA). A representative image is shown. **F-G)** Quantification of the different proteins shown in E). Tubulin was used as a loading control. Data are presented Mean ± SD unless otherwise specified, and each graph dot represents an independent biological replicate. Significant differences were assessed using Mann-Whitney U-test when comparing between two groups and considered when P<0.05. The p value is indicated in each figure.

**Supplementary Figure 5:**
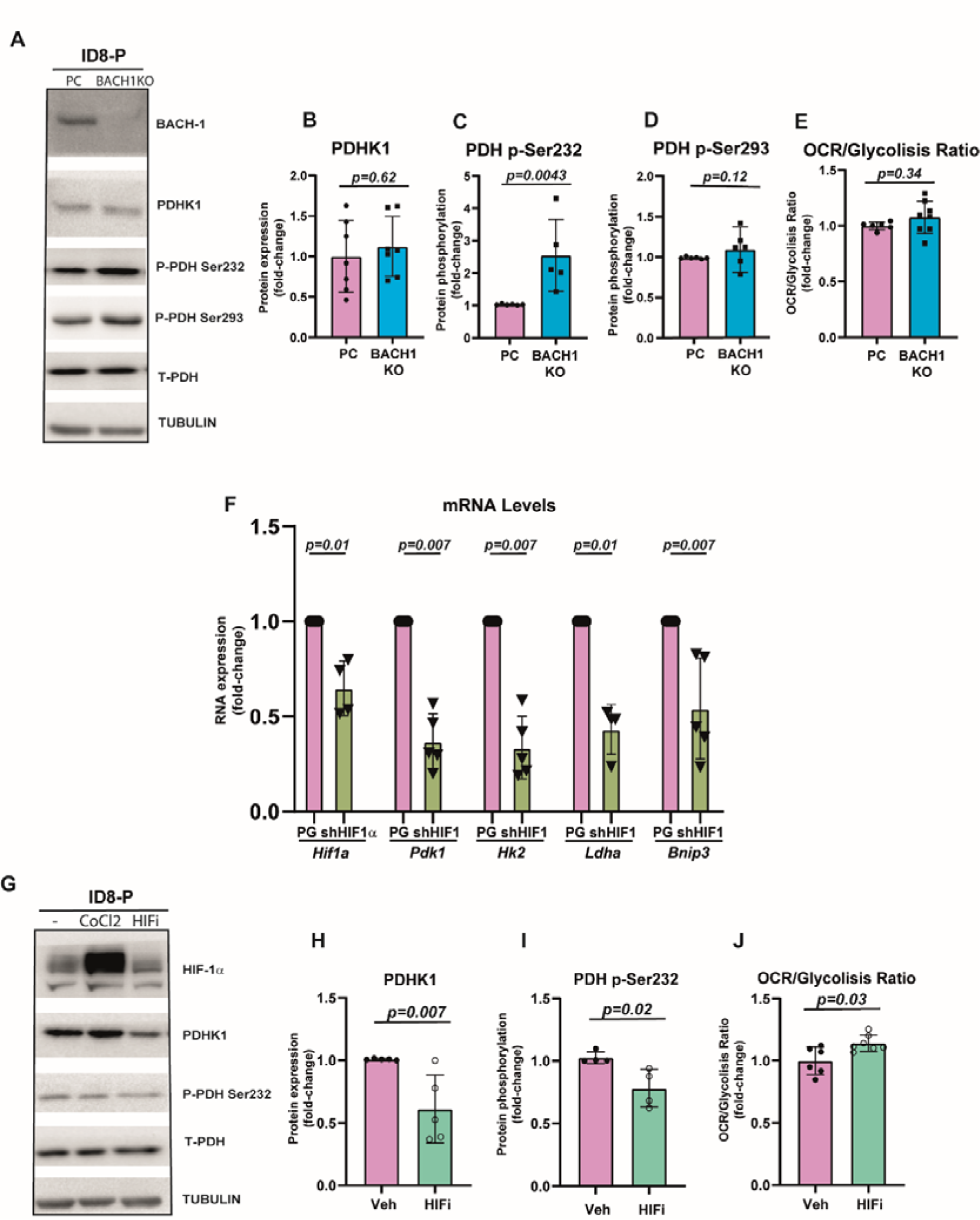
Effect of BACH1 and HIF-1. α **inhibition in ID8-P cells. A)** Western blot assessing the expression of PDHK1, the phosphorylation of serine 232 and 293 of PDH, total PDH and tubulin (as a loading control) in control (PC) and *Bach1* KO cells (BACH1 KO). A representative image is shown. **B-D)** Quantification of the different proteins shown in A). Tubulin was used as a loading control. **E)** OCR/Glycolysis ratio is shown in control (PC) and *Bach1* KO cells (BACH1 KO). **F)** mRNA levels of *Hif1a* gene and several read outs of HIF-1 activity (*Pdk1*, *Hk2*, *Ldha*, *Bnip3* genes) in control cells (ID8 P53-GFP, PG) and *Hif1a* knockdown cells (shHIF1α). *Actin* mRNA levels was used as control. **G)** Western blot assessing the expression of PDHK1, the phosphorylation in serine 293 of the PDH, total PDH and tubulin (as a loading control) in ID8-*Trp53* KO cells (ID8-P) incubated with vehicle (DMSO), cobalt chloride 200µM (CoCl2) (as a positive control of HIF-1α stimulation), or PX-478 (HIFi) 50µM for 24 hours. A representative image is shown. **H-I)** Quantification of the different proteins shown in G). Tubulin was used as a loading control. **J)** OCR/Glycolysis ratio is shown in ID8-*Trp53* KO cells (ID8-P) cells previously treated for 24h with vehicle (DMSO) PX-478 10µM (HIFi) for 24 hours. Data are presented Mean ± SD unless otherwise specified, and each graph dot represents an independent biological replicate. Significant differences were assessed using Mann-Whitney U-test when comparing between two groups and considered when P<0.05. The p value is indicated in each figure.

**Supplementary Figure 6:**
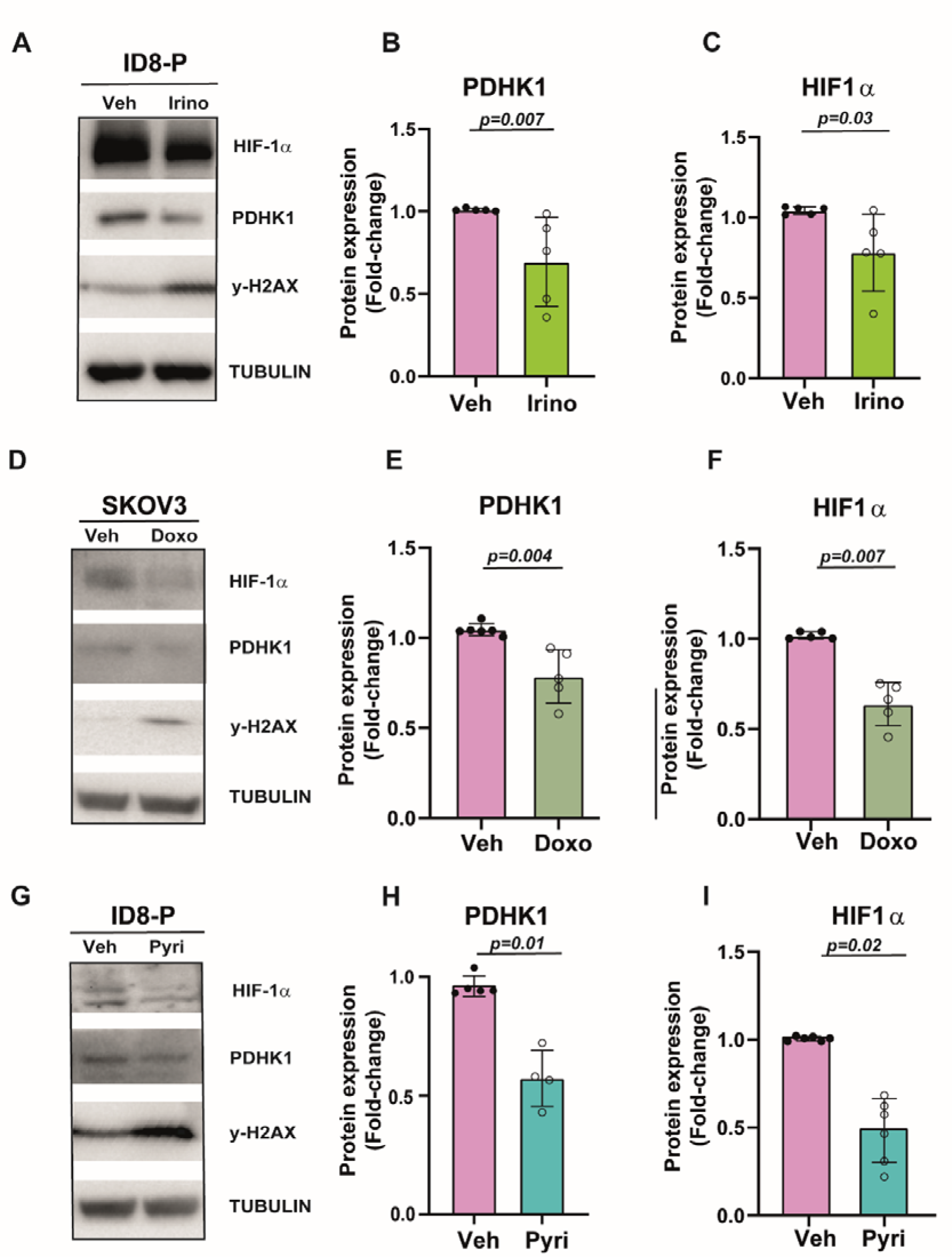
**Acute DNA damage produces a decrease of PDHK1 and HIF- 1**α **in other HGSOC models. A)** Western blot assessing the expression of PDHK1, HIF- 1α, y-H2AX and Tubulin (as a loading control) in ID8-*Trp53* KO cells (ID8-P), pretreated with vehicle (DMSO) or Irinotecan 50µM (Irino) for 24 hours. A representative image is shown. **B-C)** Quantification of PDHK1 and HIF-1α shown in A). Tubulin was used as a loading control. **D)** Western blot assessing the expression of PDHK1, HIF-1α, y-H2AX and Tubulin (as a loading control) in SKOV3N cells, pretreated with vehicle (DMSO) or Doxorubicin 2µM (Doxo) for 24 hours. A representative image is shown. **E-F)** Quantification of PDHK1 and HIF-1α shown in D). Tubulin was used as a loading control. **G)** Western blot assessing the expression of PDHK1, HIF-1α, y-H2AX and Tubulin (as a loading control) in ID8-*Trp53* KO cells (ID8-P), pretreated with vehicle (DMSO) or Pyridostatin 10µM (Pyri) for 24 hours. A representative image is shown. **H-I)** Quantification of PDHK1, HIF-1α shown in G). Tubulin was used as a loading control. Data are presented Mean ± SD unless otherwise specified, and each graph dot represents an independent biological replicate. Significant differences were assessed using Mann-Whitney U-test when comparing between two groups and considered when P<0.05. The p value is indicated in each figure.

**Supplementary Figure 7:**
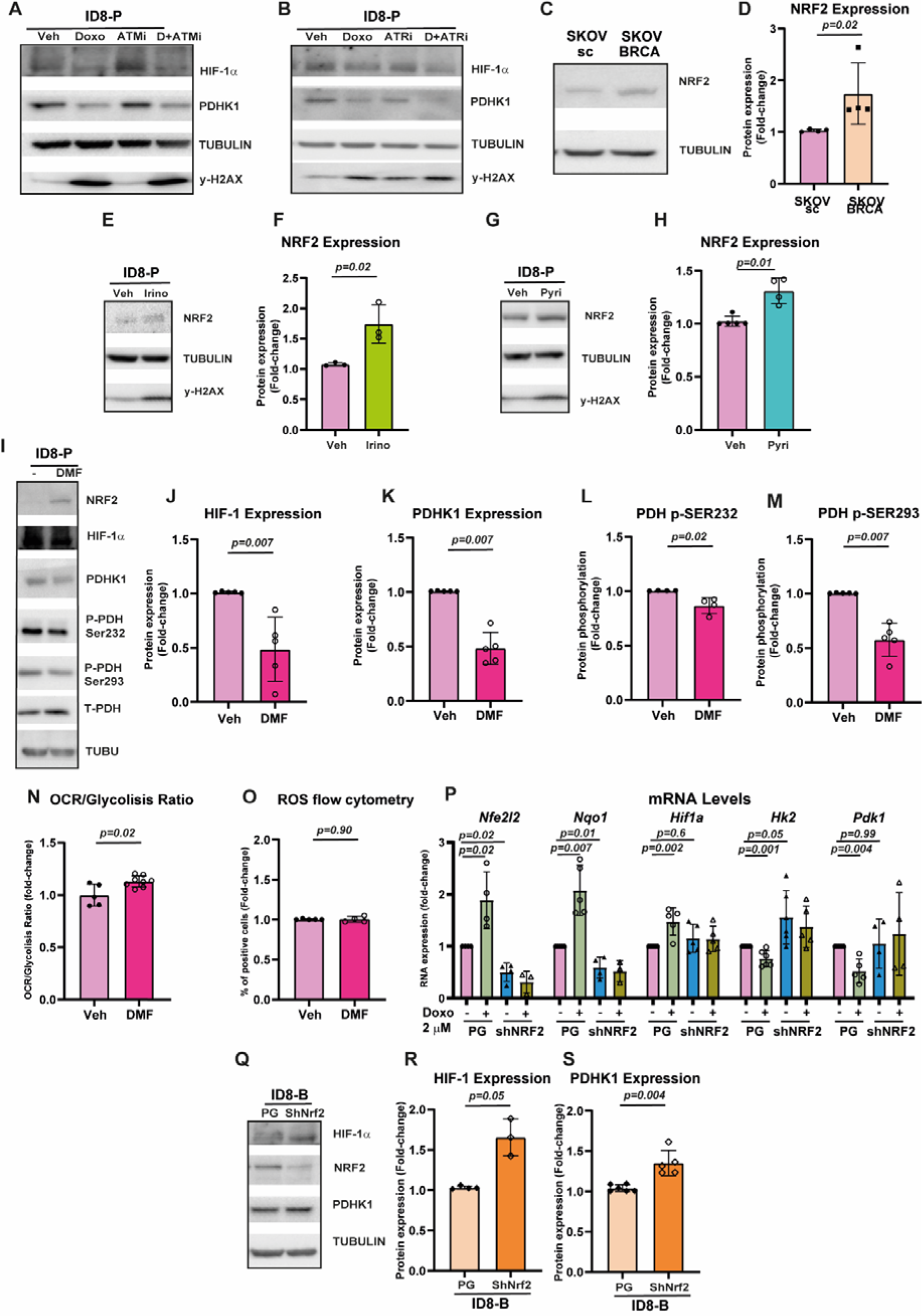
NRF2 activation is necessary to promote the metabolic shift. A) Western blot assessing the expression of PDHK1, HIF-1α, y-H2AX and Tubulin (as a loading control) in ID8-*Trp53* KO cells (ID8-P), pretreated with vehicle (DMSO), Doxorubicin 2µM (Doxo), AZD1390 20nM (ATMi) or the combination of both for 24 hours. A representative image is shown. **B)** Western blot assessing the expression of PDHK1, HIF-1α, y-H2AX and Tubulin (as a loading control) in ID8-*Trp53* KO cells (ID8-P), pretreated with vehicle (DMSO), Doxorubicin 2µM (Doxo), BAY-1895344 100nM (ATRi) or the combination of both for 24 hours. A representative image is shown. **C)** Western blot assessing the expression of NRF2 and tubulin (as a loading control) in control SKOV3N cells (SKOV-sc) and SKOV3 *BRCA2* KO cells (SKOV-BRCA). A representative image is shown. **D)** Quantification of the different proteins shown in C). Tubulin was used as a loading control. **E)** Western blot assessing the expression of NRF2, y-H2AX and Tubulin (as a loading control) in ID8-*Trp53* KO cells (ID8-P), pretreated with vehicle (DMSO) or Irinotecan 50µM (Irino) for 24 hours. A representative image is shown. **F)** Quantification of NRF2 shown in E). Tubulin was used as a loading control. **G)** Western blot assessing the expression of NRF2, y-H2AX and Tubulin (as a loading control) in ID8- *Trp53* KO cells (ID8-P), pretreated with vehicle (DMSO) or Pyridostatin 10µM (Pyri) for 24 hours. A representative image is shown. **H)** Quantification of NRF2 shown in G). Tubulin was used as a loading control. **I)** Western blot assessing the expression of NRF2, HIF-1α, PDHK1, the phosphorylation of serine 232 and 293 of PDH, total PDH and Tubulin (as a loading control) in ID8-*Trp53* KO cells (ID8-P), pretreated with vehicle (DMSO) or dimethyl fumarate 10µM (DMF) for 24 hours. A representative image is shown. **J-M)** Quantification of the different proteins shown in I). Tubulin was used as a loading control. **N)** OCR/Glycolysis ratio is shown. ID8-*Trp53* KO cells (ID8-P) were previously treated 24 hours with vehicle (DMSO) or dimethyl fumarate 10 µM (DMF). **O)** Quantification of cytosolic ROS in ID8-*Trp53* KO cells (ID8-P), pretreated with vehicle (DMSO) or dimethyl fumarate 10 µM (DMF) for 24 hours. **P)** mRNA levels of *Nfe2l2, Nqo1, Hif1a, Hk2, Pdk1* genes in control cells (ID8 P53-GFP, PG) and *Nfe2l2* knockdown cells (shNRF2) pretreated with vehicle (DMSO) or doxorubicin 2µM (Doxo) for 24 hours. *Actin* mRNA levels was used as control. **Q)** Western blot assessing the expression of NRF2, HIF-1α, PDHK1 and tubulin (as a loading control) in control (ID8 BRCA-tdTomato, PG) and *Nfe2l2* knockdown cells (shNRF2). A representative image is shown. **R-S)** Quantification of the different proteins shown in Q). Tubulin was used as a loading control. Data are presented Mean ± SD unless otherwise specified, and each graph dot represents an independent biological replicate. Significant differences were assessed using Mann-Whitney U-test when comparing between two groups and considered when P<0.05. The p value is indicated in each figure.

**Supplementary Figure 8:**
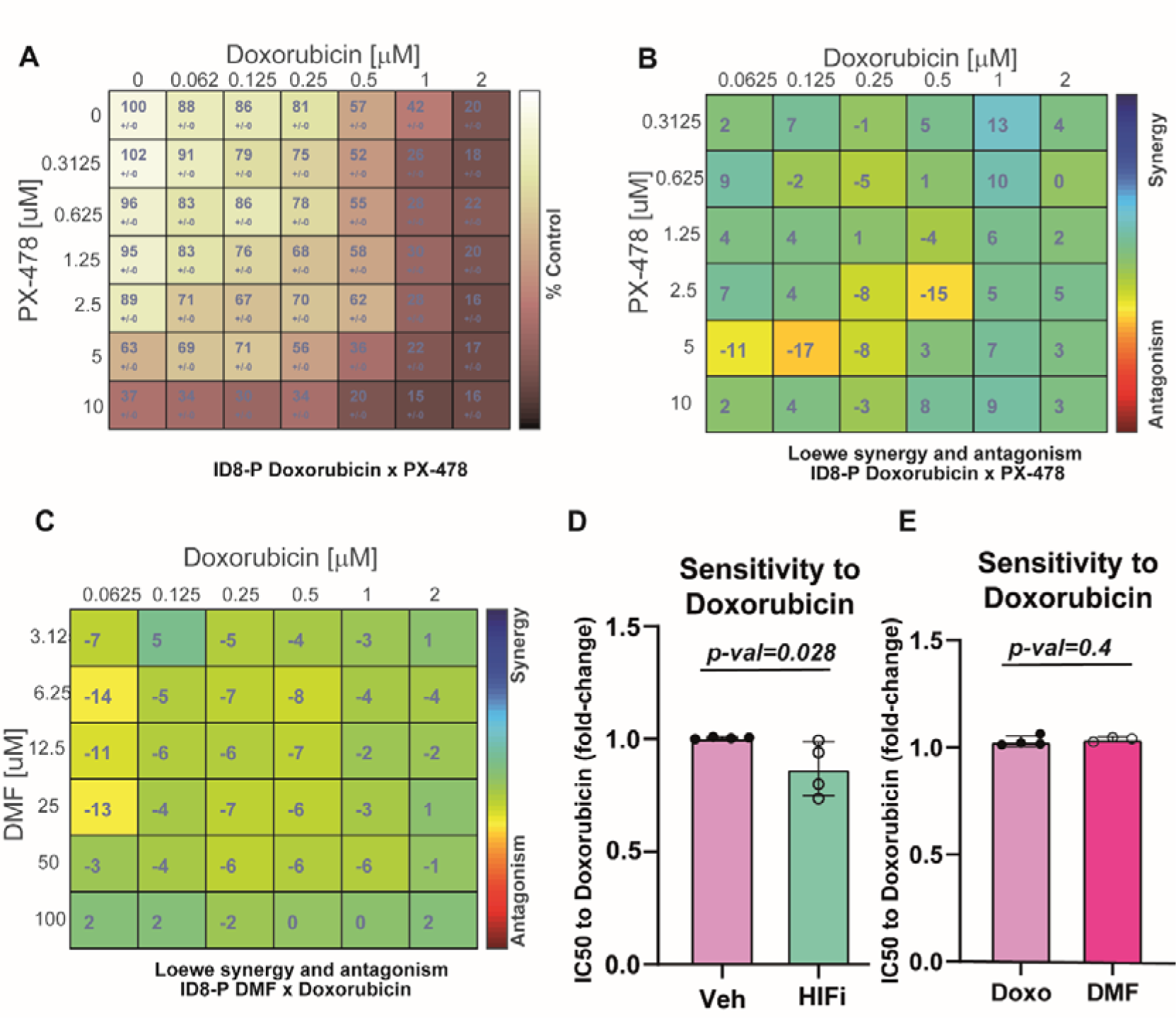
The combination of chemotherapy and HIF-1 inhibitors or NRF-2 activators does not produce a synergistic effect. A) Viability matrix of ID8-Trp53 KO cells (ID8-P) incubated with different doses of doxorubicin and PX-478. ID8-Trp53 KO cells (ID8-P) were seeded and treated with increasing doses of each drug as shown, for 72 hours **B)** Synergy matrix of ID8-Trp53 KO cells (ID8-P) incubated with different doses of doxorubicin and PX-478. **C)** Synergy matrix of ID8-Trp53 KO cells (ID8-P) incubated with different doses of doxorubicin and dimethyl fumarate (DMF). **D)** Graph showing the IC50 to doxorubicin. ID8-Trp53 KO cells (ID8-P), treated with vehicle (DMSO) or PX-478 5µM (HIFi) and then with increasing doses of doxorubicin for 72 hours. IC50 was assessed with the non-linear regression tool from GraphPad Prism 8. **E)** Graph showing the IC50 to doxorubicin. ID8-Trp53 KO cells (ID8-P), treated with vehicle (DMSO) or dimethyl fumarate 10µM (DMF) and then with increasing doses of doxorubicin for 72 hours. IC50 was assessed with the non-linear regression tool from GraphPad Prism 8. Data are presented Mean ± SD unless otherwise specified, and each graph dot represents an independent biological replicate. Significant differences were assessed using Mann-Whitney U-test when comparing between two groups and considered when P<0.05. The p value is indicated in each figure.

**Supplementary Figure 9:**
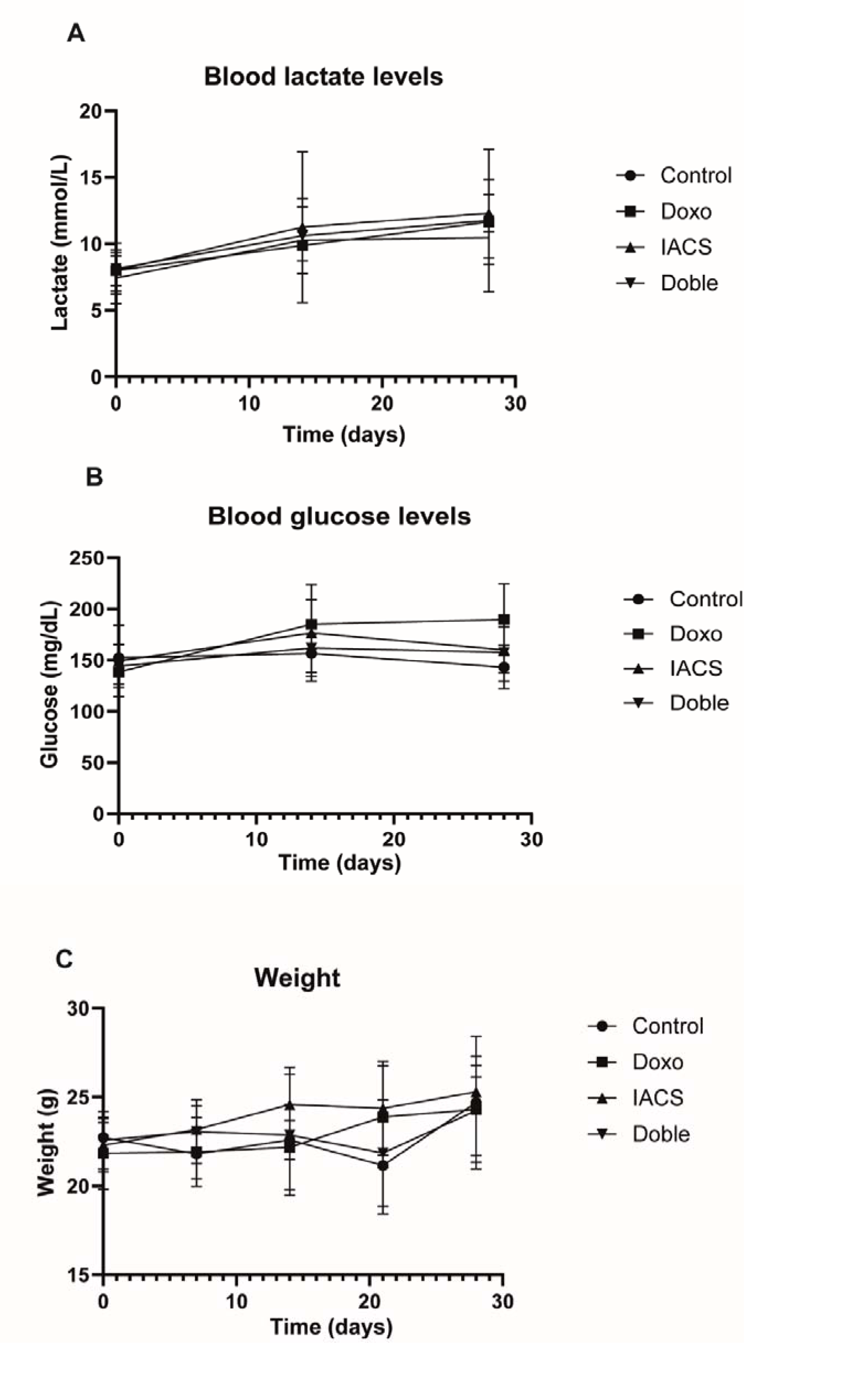
Doxorubicin and OXPHOS inhibitors do not affect basal levels of lactate, weight and glucose in vivo. A-C) Graphs showing the evolution of lactate (A), glucose (B) blood levels and weight (C) in vivo. 0 days is the day of the start of the treatment. In each graph is represented Mean ± SD unless otherwise specified.

**SUPPLEMENTARY TABLE 1:** Correlations from the Database of genomics of drug sensitivity in cancer of the Sanger Institute between the sensitivity to Dihydrorotenone and the selected ssGSEA expression. ssGSEA were selected from the canonical pathways of the curated gene sets of the GSEA and were related to DNA damage, glycolytic metabolism or oxidative metabolism. Genesets are shown if Spearman estimate was >|0.1| and FDR<0.05.

|  | ssGSEA | SPEARMAN<br>ESTIMATE | SPEARMAN<br>P-VALUE | SPEARMAN<br>FDR |
| --- | --- | --- | --- | --- |
| NEGATIVE CORRELATION<br>(MORE EXPRESSION MORE SENSITIVE) | REACTOME_RESPIRATORY_ELECTRON_TRANSPORT | -0.24629 | 1.13471E-07 | 3.81263E-06 |
|  | REACTOME_BASE_EXCISION_REPAIR | -0.24225 | 1.84249E-07 | 5.15898E-06 |
|  | REACTOME_MITOCHONDRIAL_TRANSLATION | -0.22327 | 1.60608E-06 | 2.45292E-05 |
|  | HALLMARK_OXIDATIVE_PHOSPHORYLATION | -0.22227 | 1.79135E-06 | 2.50789E-05 |
|  | PID_FANCONI_PATHWAY | -0.20977 | 6.72654E-06 | 6.6474E-05 |
|  | REACTOME_COMPLEX_I_BIOGENESIS | -0.19847 | 2.08291E-05 | 0.000159059 |
|  | KEGG_CITRATE_CYCLE_TCA_CYCLE | -0.18679 | 6.28624E-05 | 0.000391427 |
|  | REACTOME_DNA_DOUBLE_STRAND_BREAK_REPAIR | -0.18398 | 8.11947E-05 | 0.00047037 |
|  | KEGG_MISMATCH_REPAIR | -0.1783 | 0.000134764 | 0.000665892 |
|  | KEGG_OXIDATIVE_PHOSPHORYLATION | -0.17758 | 0.000143572 | 0.000670005 |
|  | BIOCARTA_ATRBRCA_PATHWAY | -0.17669 | 0.000155051 | 0.000704015 |
|  | PID_ATM_PATHWAY | -0.17577 | 0.000167989 | 0.000723644 |
|  | GOMF_DNA_TOPOISOMERASE_ACTIVITY | -0.17486 | 0.000181729 | 0.000744644 |
|  | REACTOME_TRANSCRIPTIONAL_ACTIVATION_OF_MITOCHON<br>DRIAL_BIOGENESIS | -0.16896 | 0.000299935 | 0.001095416 |
|  | PID_ATR_PATHWAY | -0.16212 | 0.000525292 | 0.001604528 |
|  | PID_HIF1A_PATHWAY | -0.14554 | 0.001871851 | 0.004557551 |
|  | REACTOME_DNA_DOUBLE_STRAND_BREAK_RESPONSE | -0.14336 | 0.00219267 | 0.005084439 |
|  | REACTOME_MITOCHONDRIAL_BIOGENESIS | -0.14326 | 0.00220931 | 0.005084439 |
|  | KEGG_HOMOLOGOUS_RECOMBINATION | -0.13661 | 0.00352789 | 0.007598532 |
|  | REACTOME_THE_CITRIC_ACID_TCA_CYCLE_AND_RESPIRATOR<br>Y_ELECTRON_TRANSPORT | -0.12555 | 0.007365087 | 0.013748162 |
|  | REACTOME_GLUCOSE_METABOLISM | -0.11387 | 0.015127556 | 0.025162667 |
|  | REACTOME_NONHOMOLOGOUS_END_JOINING_NHEJ | -0.10665 | 0.02294112 | 0.03535879 |
|  | REACTOME_CITRIC_ACID_CYCLE_TCA_CYCLE | -0.10187 | 0.029840176 | 0.044364155 |
| POSITIVE CORRELATION<br>(MORE EXPRESSION MORE<br>RESISTANT) | REACTOME_REGULATION_OF_GLYCOLYSIS_BY_FRUCTOSE_2_<br>6_BISPHOSPHATE_METABOLISM | 0.107991 | 0.02126596 | 0.033080382 |
|  | REACTOME_REGULATION_OF_GENE_EXPRESSION_BY_HYPOXI<br>A_INDUCIBLE_FACTOR | 0.108361 | 0.020824455 | 0.032696341 |
|  | BIOCARTA_HIF_PATHWAY | 0.110226 | 0.018717426 | 0.029947882 |
|  | BIOCARTA_P53HYPOXIA_PATHWAY | 0.171479 | 0.000242676 | 0.000948131 |
|  | REACTOME_PYRUVATE_METABOLISM | 0.212832 | 4.89731E-06 | 5.3102E-05 |
|  | REACTOME_REGULATION_OF_HMOX1_EXPRESSION_AND_AC<br>TIVITY | 0.216538 | 3.31654E-06 | 3.97985E-05 |
|  | REACTOME_PTK6_PROMOTES_HIF1A_STABILIZATION | 0.231468 | 6.44277E-07 | 1.20265E-05 |
|  | HALLMARK_GLYCOLYSIS | 0.300967 | 6.95398E-11 | 5.84134E-09 |
|  | HALLMARK_HYPOXIA | 0.338949 | 1.46327E-13 | 2.45829E-11 |

**SUPPLEMENTARY TABLE 2:** Mutations in *P53* gene and genes related to homologous recombination repair in the used cell lines with the indicated reference.

| MUTATIONS | P53 | BRCA1 | BRCA2 | RAD51 | ATM |
| --- | --- | --- | --- | --- | --- |
| ID8-P | X<sup>30</sup> |  |  |  |  |
| ID8-B | X<sup>30</sup> |  | X<sup>30</sup> |  |  |
| SKOV | X<sup>59</sup> |  |  |  |  |
| A2780 |  |  |  |  | X<sup>60</sup> |
| OVCAR4 | X<sup>59</sup> |  |  |  |  |
| 22RV1 | X<sup>61</sup> |  | X<sup>61</sup> |  |  |
| NP P53 | X<sup>62</sup> |  |  |  |  |
| DU145 | X<sup>59,61</sup> |  |  |  |  |
| T-47 D | X<sup>59</sup> |  |  |  |  |

**SUPPLEMENTARY TABLE 3:** TGCA Study Abbreviations used in the present work.

| ABBREVIATION | COMPLETE NAME |
| --- | --- |
| ACC | Adrenocortical carcinoma |
| BLCA | Bladder Urothelial Carcinoma |
| BRCA | Breast invasive carcinoma |
| CESC | Cervical squamous cell carcinoma and endocervical adenocarcinoma |
| CHOL | Cholangiocarcinoma |
| COAD | Colon adenocarcinoma |
| DLBC | Lymphoid Neoplasm Diffuse Large B-cell Lymphoma |
| ESCA | Oesophageal carcinoma |
| GBM | Glioblastoma multiforme |
| HNSC | Head and Neck squamous cell carcinoma |
| KICH | Kidney Chromophobe |
| KIRC | Kidney renal clear cell carcinoma |
| KIRP | Kidney renal papillary cell carcinoma |
| LGG | Brain Lower Grade Glioma |
| LIHC | Liver hepatocellular carcinoma |
| LUAD | Lung adenocarcinoma |
| LUSC | Lung squamous cell carcinoma |
| MESO | Mesothelioma |
| OV | Ovarian serous cystadenocarcinoma |
| PAAD | Pancreatic adenocarcinoma |
| PCPG | Pheochromocytoma and Paraganglioma |
| PRAD | Prostate adenocarcinoma |
| READ | Rectum adenocarcinoma |
| SARC | Sarcoma |
| SKCM | Skin Cutaneous Melanoma |
| STAD | Stomach adenocarcinoma |
| TGCT | Testicular Germ Cell Tumours |
| THCA | Thyroid carcinoma |
| THYM | Thymoma |
| UCEC | Uterine Corpus Endometrial Carcinoma |
| UCS | Uterine Carcinosarcoma |
| UVM | Uveal Melanoma |

